# *Nf1* deficiency accelerates mammary development and promotes luminal-basal plasticity

**DOI:** 10.1101/2022.12.22.520633

**Authors:** Elizabeth A. Tovar, Menusha Arumugam, Curt J. Essenburg, Patrick S. Dischinger, Jamie L. Grit, Megan E. Callaghan, Rachael T.C. Sheridan, Lisa Turner, Corinne R. Esquibel, Kristin Feenstra, Zachary B. Madaj, Ian Beddows, Carrie R. Graveel, Matthew R. Steensma

## Abstract

The tumor suppressor *NF1* is a critical driver of sporadic breast cancer and NF-related breast cancers. We utilized distinct *Nf1*-deficient immunocompetent rat models to investigate *Nf1* function in mammary development and homeostasis. Here we demonstrate that *Nf1* deficiency dramatically accelerates mammary morphogenesis, alters TEB cell organization, and proliferation. Notably, we observed a shift in luminal-basal epithelial lineage commitment within *Nf1*-deficient lines with early tumor onset. In addition, we detected subpopulations of hybrid EMT cells (Ecad^+^/CK14^+^) within the invasive edge and stroma of *Nf1*-deficient tumors. *Nf1* deficiency restricted luminal progenitor potential and resulted in gene expression changes associated with decreased cell adhesion and increased EMT signatures. Together our findings support a model in which *Nf1* loss of function results in lineage plasticity throughout mammary morphogenesis and promotes EMT-mediated invasion. This study reveals a previously unknown role for the tumor suppressor neurofibromin in mammary homeostasis and phenotypic plasticity during breast cancer progression.

## INTRODUCTION

Recent studies have revealed that the tumor suppressor *NF1* is a genetic driver of sporadic breast cancer and endocrine resistance.^1–6^ *NF1* is the key negative regulatory gene of the RAS pathway and the *NF1* gene product, neurofibromin, stimulates intrinsic RAS GTPase activity through binding to the GAP-related domain (GRD).^7^ Loss-of-function or genomic alterations in *NF1* result in dysregulated RAS and activation of the RAS-ERK signaling pathway. Our limited understanding of *NF1* is primarily based on studies of the tumor predisposition syndrome Neurofibromatosis Type 1 (NF). NF is caused by mutation of the *NF1* gene and is the most common single-gene disorder, affecting 1 in 3,000 live births.^8^,^9^ Even though NF is most commonly associated with benign and malignant neurofibromas, both women and men with NF have a significantly increased risk of dying from breast cancer.^10–13^ Two recent Finnish studies determined that in women with NF <40 years old have a 10-fold increased risk of breast cancer compared to the normal population.^14^ Moreover, NF-related breast cancers are associated with adverse prognostic factors and decreased overall survival compared to sporadic breast cancer.^10^ The correlation of *NF1* loss-of-function in both NF-related and sporadic breast cancers indicates that *NF1* may play a critical role in mammary homeostasis.

The breast is a highly dynamic organ composed of two major specialized epithelial cell types including 1) luminal cells that line the breast ducts and have secretory functions as well as 2) the basal/myoepithelial cells that have contractile functions.^15^ Investigations into mammary epithelial cell morphogenesis and differentiation have exposed the origins of tumor initiating cells and their relationship to mammary stem and progenitor cells. Currently, there is a lack of understanding on the role of *NF1* in mammary gland differentiation and breast cancer initiation. Much of our understanding of the functional loss of *NF1* and tumorigenesis come from studies of genetically engineered mouse models which do not fully recapitulate human disease, or other phenotypic aspects of *NF1* deficiency.^16–19^ *Nf1*-deficient models have been valuable in defining how *Nf1* promotes neurofibroma development and malignant progression; yet breast cancer is not an observed phenotype in any *Nf1*-deficient animal models. To develop a more comprehensive model of *Nf1* deficiency in breast cancer, we utilized CRISPR-Cas9 gene editing to create germline *Nf1* indels in immunocompetent Sprague Dawley rats. To impact RAS regulation, we generated several distinct *Nf1* indels in exon 20 of the GRD that resulted in either in-frame or premature stop deletions.^20^ The *Nf1* in-frame (*Nf1^IF^*) animals were the result of a 54-bp deletion in exon 20, which altered the GRD. The premature stop (*Nf1^PS^*) indel resulted from an 8-bp deletion in exon 20 (Figure 1A); consequently, all of the functional domains beyond exon 20, including most of the GRD, are not expressed. The *Nf1^PS-21^* line is derived from a *Nf1^IF^* indel (57-bp deletion in exon 20) that has a distinct mRNA isoform with an additional 140-bp deletion in exon 21. This exon 21 deletion mRNA was exclusive to the *Nf1^PS-21^* line and resulted in a premature stop that impacts the GRD domain and other downstream functional domains.^20^ All of these indels model “*Nf1* deficiency” and induced highly penetrant, aggressive mammary adenocarcinomas in female and male rats. Immunostaining demonstrated that *Nf1*-deficient tumors were ER+/PR+.^20^ Analysis of survival revealed that *Nf1^PS^* and *Nf1^PS-21^* animals developed more aggressive tumors compared to *Nf1^IF^* rats, which was not unexpected based on the predicted partial functionality of the *Nf1* indels (Figure 1A). Our data strongly extend these findings as we show that both mutation type (IF and PS) and splice variation lead to more aggressive disease. The phenotypes of our *Nf1* rat model and the identification of novel *Nf1* mRNA and protein isoforms underscore our limited understanding of *NF1* expression and function.

**Figure 1:**
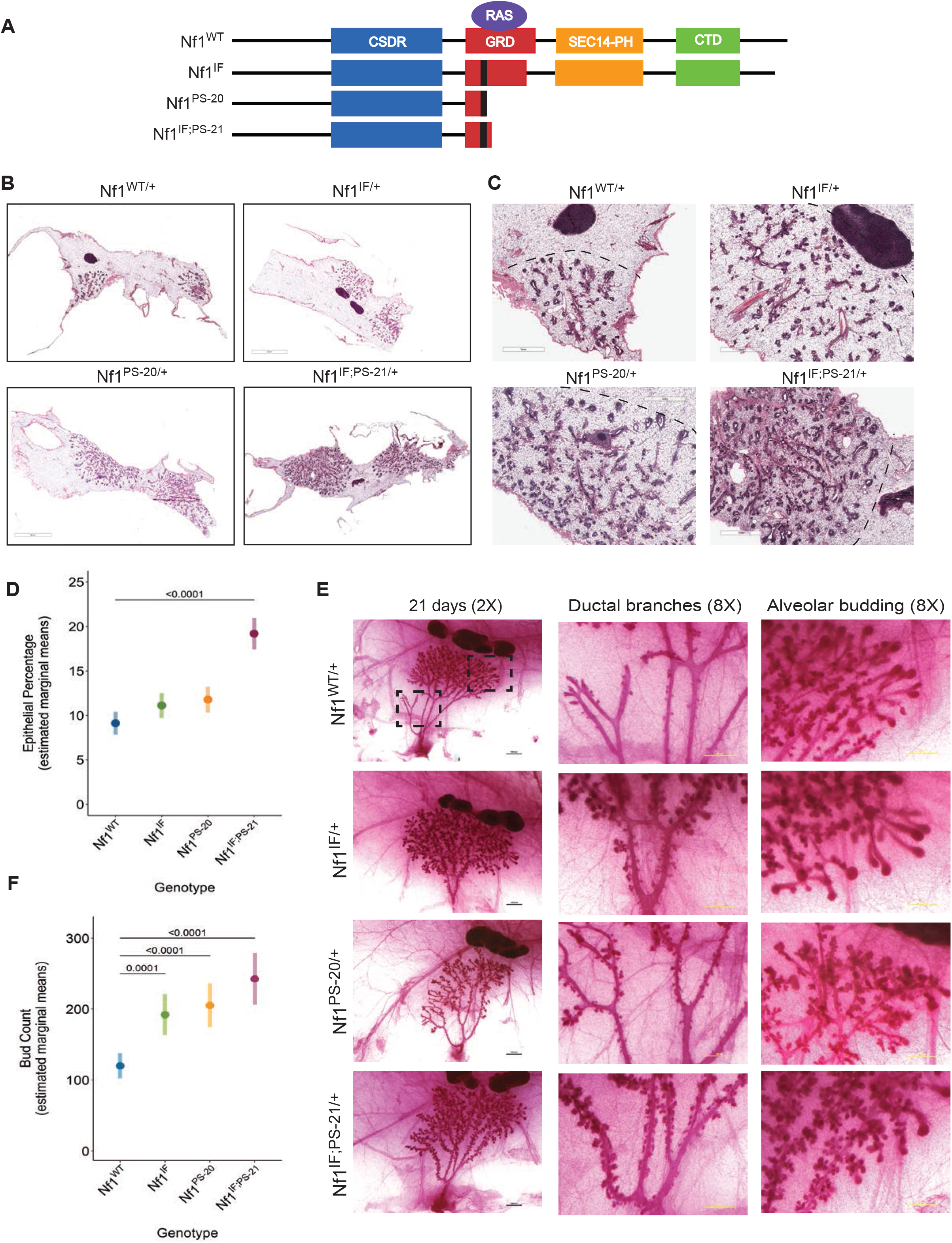
*Nf1* deficiency promotes aberrant ductal outgrowth and alveolar expansion during mammary morphogenesis. A) Schematic of wild-type and mutant neurofibromin in Sprague Dawley rats. The black bar within the GRD denotes indel location from CRISPR editing. Immunohistochemistry H&E sections of wild-type and *Nf1*-deficient glands at B) 5x [scale bar = 5mm] and C) 20x magnification [scale bar = 1mm]. The dashed black line indicates the tip of the ductal tree in relation to the lymph node. D) 95% confidence intervals for mean percentage of epithelial cells per genotype (n=4) quantified from H&E images. E) Carmine stains of whole mammary fat pads from 21-day old wild-type and *Nf1*-deficient rats showing the entire ductal tree at 20x (first column, scale bar = 1000 μm), ductal branches at 80x (second column, scale bar = 500 μm), and alveolar buds/TEBs (last column) at 80x magnification. F) 95% confidence intervals for mean total alveolar and TEB counts quantified per genotype (WT vs. IF p = 0.0001, WT vs. PS-20 or IF;PS-21 p<0.0001; n=5) from 80x carmine images.

To interrogate the impact of somatic *NF1* loss in breast cancer, we previously performed an analysis of the METABRIC human breast cancer dataset.^21^ Similar to what has been observed in other breast cancer datasets, 2.1% of tumors harbored *NF1* mutations.^1^,^2^ Surprisingly, genomic alterations in *NF1* were identified in 27% of sporadic breast cancers, with copy number alterations (CNAs) being the dominant alteration (25%). A survival analysis revealed that patients with *NF1* shallow deletions are more likely to have basal-like breast cancer, less likely to have luminal A breast cancers, and have a poorer 10 year survival. Gene correlation network analysis revealed a strong association of *NF1* shallow deletions in ER+ and ER-tumor subsets and with endocrine-resistance genes. These results indicate that *NF1* genomic alterations in breast cancer are more common than previously thought and are present in ER+ and ER-tumors.

To determine how *NF1* deficiency promotes tumor initiation in the breast, it is essential to understand the mechanistic role *NF1* has in normal mammary development and homeostasis. Here we show *Nf1* deficiency accelerates mammary morphogenesis and alters mammary homeostasis at the earliest stages. By optimizing established protocols for isolation and characterization of rat mammary epithelial populations, we dissected the impact of *Nf1* deficiency on the mammary epithelial hierarchy. Collectively, these results revealed a shift in luminal to basal lineage commitment within the *Nf1*-deficient lines that have the earliest tumor onset. *Nf1* deficiency in both rat mammary tissues and human MCF10A breast cells resulted in epithelial-to-mesenchymal (EMT) signatures and the presence of subpopulations with luminal-basal plasticity. Therefore, our findings demonstrate that *Nf1* loss of function promotes a shift in lineage commitment within the mammary epithelial hierarchy which correlates with tumor incidence. The results of this study highlight a previously unknown role for the tumor suppressor neurofibromin in maintaining mammary homeostasis during development and may indicate the basis for higher breast cancer risk in individuals with NF and the poor outcomes within *NF1*-related sporadic breast cancers.

## RESULTS

### *Nf1* deficiency promotes accelerated mammary morphogenesis

To understand how *Nf1* deficiency promotes tumor initiation in the breast, we explored the impact of *Nf1* indels and neurofibromin function on mammary morphogenesis. To start, we examined several timepoints of postnatal mammary development in our *Nf1*-deficient rat models (Figure 1A). Mammary gland whole mounts from the 4^th^ mammary fat pad were analyzed at postnatal day 21 which represents the start of puberty. At this timepoint, the development and expansion of both the 4^th^ and 5^th^ ductal trees into the mammary fat pad can be observed (Figure 1B). Examination of the ductal outgrowth patterns in 21-day old females demonstrated a substantial increase in ductal expansion in all the *Nf1*-deficient lines compared to *Nf1^WT/+^* mammary pads (Figures 1B-C). At postnatal day (PND) 21, the *Nf1^WT/+^* ductal outgrowths had elongated to approximately 2/3 of the distance from the nipple to the lymph node (LN). In contrast, both *Nf1^IF/+^* and *Nf1^PS-20/+^* females demonstrated accelerated ductal expansion that reached or surpassed the 4^th^ mammary gland LN (Figure 1B-C). Even though the *Nf1^IF;PS-21/+^* ductal expansion did not extend longitudinally to the extent of that in *Nf1^IF/+^* and *Nf1^PS-20/+^* ductal outgrowths, the *Nf1^IF;PS-21/+^* pads showed expansive ductal filling throughout the fat pad compared to the *Nf1^WT/+^* and other *Nf1*-deficient lines (Figure 1B-C). Automated quantitation revealed a significant increase in the percent of epithelial cells present in the 4^th^ mammary fat pads in all the *Nf1*-deficient lines compared to *Nf1^WT/+^* glands (*p* < 0.0001; Figure 1D). These results imply that *NF1* plays a role in regulating mammary gland development.

To investigate the impact of *Nf1* deficiency on ductal elongation, we evaluated both the structural patterns of ductal branching and terminal end buds (TEBs). TEBs are bulbous formations at the tips of developing ducts that are completely unique to the peripubertal mammary gland and believed to contain mammary stem cells (MaSCs). The MaSCs within the TEB are responsible for the production of the mature cell types that result in elongation of the subtending duct.^22^ Carmine staining on wholemounts of the mammary glands demonstrated striking differences in both the ductal and TEB structures in the *Nf1*-deficient lines (Figure 1E). At 21 days, the primary ductal branches had no signs of lateral ductal budding in the *Nf1^WT/+^* mammary pad, whereas there was profuse budding on subtending primary ducts in the *Nf1*-deficient lines (Figure 1E, middle panel). The TEBs in the *Nf1^IF/+^* line were slightly larger than those in the *Nf1^WT/+^* mammary gland, and the TEBs in the *Nf1^PS-20/+^* and *Nf1^IF;PS-21/+^* lines were a mix of large TEBs and small, grape-like clusters of aggregated buds pointed in multiple directions, indicating disorganized directional formation coupled with aberrant cell growth (Figure 1E, right panel). Blinded quantification of the total number of buds, both TEBs and lateral, revealed an increased number of buds in the *Nf1*-deficient lines compared to *Nf1^WT/+^* (Figure 1F). To get a better understanding of the timeline in which *Nf1* influences mammary development we also assessed glands at PND 8 and 33 (Figure S1). Accelerated ductal outgrowth and aberrant alveolar budding is present at PND 8 in *Nf1*-deficient mammary glands. At PND 33 the *Nf1*-deficient mammary glands have pervaded throughout the fat pad more extensively than *Nf1^WT/+^* glands. These findings demonstrate that *Nf1* deficiency results in accelerated mammary development that is mitigated by the level of *Nf1* functional loss, such that *Nf1^PS-20/+^* and *Nf1^IF;PS-21/+^* mammary glands have more extensive aberrant ductal growth compared to *Nf1^IF/+^* mammary glands at these early stages of development.

### *Nf1* deficiency alters TEB cell organization and proliferation

To gain further insight into the impact of *Nf1* function on pubertal mammary development, we focused on the cells comprising the 21 day TEB. The inner mass of a TEB is made of body cells, which are presumed to be luminal ductal and alveolar progenitors. The single-cell layer externally lining the TEB are composed of cap cells, which give rise to myoepithelial cells. Cap cells have been shown to have regenerative stem cell capabilities in transplantation assays, but other studies have contradicted these results, and whether cap cells are MaSCs or restricted progenitors is still debated.^23^ A combination of epithelial and stromal signals are essential for maintaining TEB unidirectional and non-overlapping growth.^24^–^26^ High magnification images showed that *Nf1^IF/+^* TEBs are nearly indistinguishable from *Nf1^WT/+^* TEBs, except for the presence of larger TEBs within close proximity (Figure 2A). The TEBs from *Nf1^PS-20/+^* and *Nf1^IF;PS-21/+^* lines were considerably distorted compared to *Nf1^WT/+^* glands. Notably, TEBs from *Nf1^PS-20/+^* glands had a thickened body cell layer while *Nf1^IF;PS-21/+^* TEBs were an amalgamation of normal and completely disorganized buds that resembled the terminal ductal-lobular units present in adult human glands.^27^ Note that similar tubuloalveoar formations are not observed in rats until after post-natal day 46, which is beyond the 21 day timepoint shown in Figure 2.^28^ These results suggest that *Nf1* deficiency perturbs TEB formation and consequently ductal outgrowth in mammary development.

**Figure 2:**
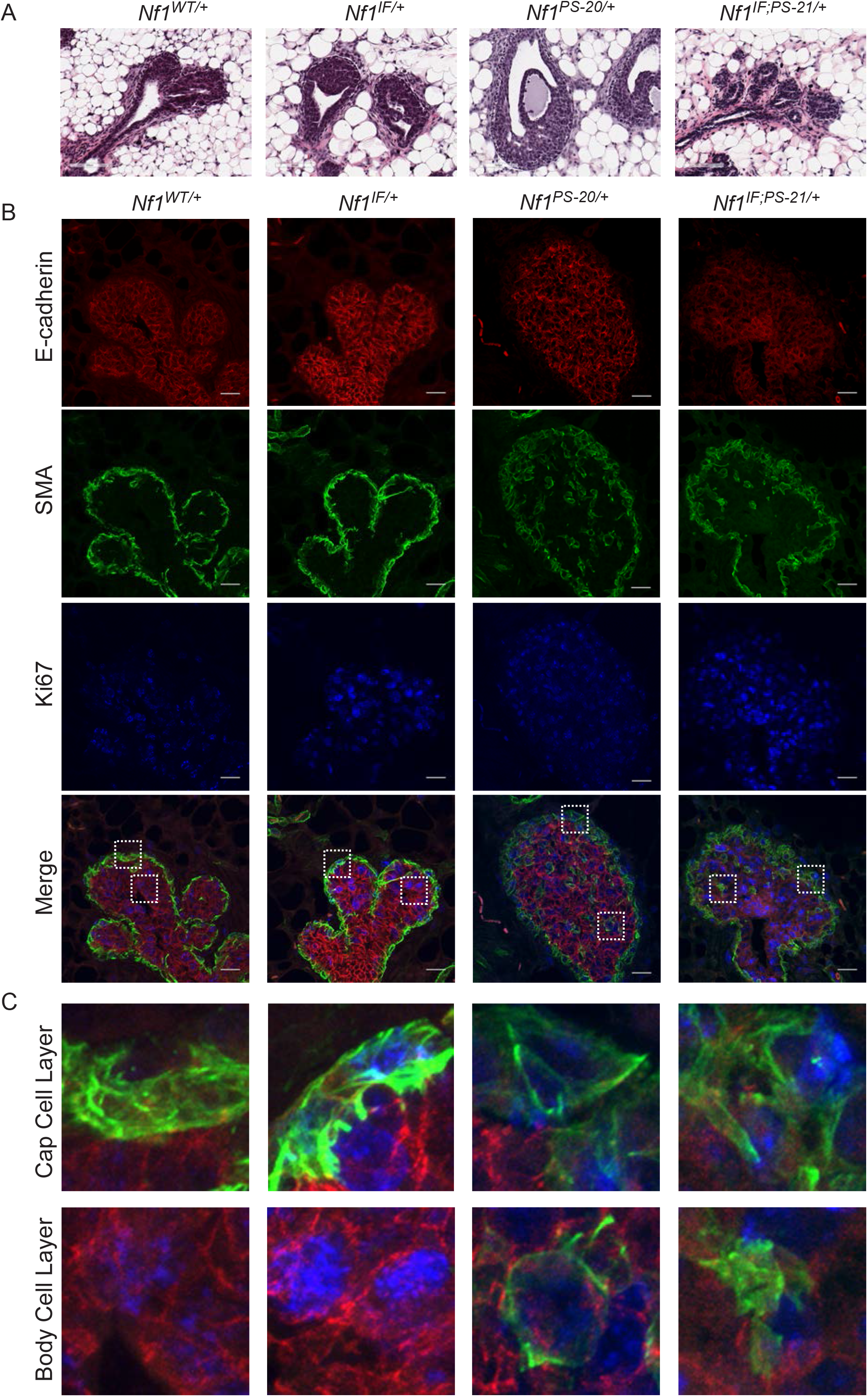
*Nf1* indels impact mammary morphogenesis and TEB formation. Immunohistochemistry H&E of TEBs from 21-day old female rats. Images were taken at 200x magnification, scale bar = 100 μm. B) Immunofluorescent confocal images of TEBs from 21-day old female rats immunostained with E-cadherin (red), SMA (green), and Ki67 (blue). Images were acquired at 600x magnification, scale bar = 21 μm. C) Insets from part B ‘Merge’ column. Upper row shows cells in the CAP cell layer and the bottom row shows cells in the body cell compartment.

To elucidate how *Nf1* deficiency alters TEB structure and cell proliferation, we immunostained mammary gland sections from PND 21 females with distinct, well-defined lineage markers: smooth muscle actin (SMA, green) for basal/myoepithelial/cap cells and E-cadherin (red) for luminal epithelial/body cells, combined with Ki67 to mark proliferating cells (blue) (Figure 2B-C). ^23^ Compared to *Nf1^WT/+^*, the overall structure of *Nf1^IF/+^* TEBs was comparable with similar expression patterns of E-cadherin in the TEB body and SMA at the TEB periphery (Figure 2B). In contrast, *Nf1^PS-20/+^* and *Nf1^IF;PS-21/+^* TEBs had diffuse and disorganized E-cadherin and SMA expression throughout the TEB. As shown in the magnified images of the *Nf1^WT/+^* TEB in Figure 2C, body cells are enveloped by a contiguous single-cell layer of cap cells. The rare cap cells that are found in the body cell layer are hypothesized to be left behind after TEB outgrowth ^29^ and undergo cell death.^23,30^ In *Nf1^PS-20/+^* TEBs, we observed decreased and diffuse SMA staining at the periphery and the presence of SMA+/Ecad+ cells within the body cell layer. A similar, but more pronounced phenotype was observed in *Nf1^IF;PS-21/+^* glands, where E-cadherin expression was diminished throughout the TEB body and numerous SMA/Ecad+ cells were observed (Figure 2B-C). Evaluation of cap cell proliferation in the *Nf1*-deficient TEBs revealed a marked increase in Ki67 staining intensity in cap cell nuclei in all the *Nf1*-deficient lines compared to *Nf1^WT/+^,* yet the strongest Ki67 staining was present in *Nf1^IF/+^* and *Nf1^IF;PS-21/+^* TEBs (Figure 2B). We also detected Ki67 staining in the cap cells within the TEB body with weak E-cadherin staining (Figure 2C). The increased proliferation was verified by immunohistochemical staining of Ki67 in cross-sections of the developing mammary glands (Figure S2). Together the presence of SMA+/Ecad+ cells, decreased E-cadherin expression, and altered cap cell localization suggests that *Nf1* deficiency impacts epithelial differentiation within the developing mammary gland.

### *Nf1* deficiency induces a shift in luminal to basal epithelial populations

Methods to characterize the epithelial hierarchy have been well established in the human and mouse mammary glands.^31^ To delineate whether *Nf1* deficiency alters the mammary epithelial hierarchy, we modified and optimized these methods for rat tissues.^32^ We isolated mammary epithelial cells (MECs) by enzymatic digestion of the 4^th^ mammary fat pads from 22 and 43 day old female rats, and then subjected the samples to FACS (fluorescence-activated cell sorting). Briefly, endothelial and hematopoietic cells were depleted from freshly digested samples using the lineage markers CD31 and CD45 respectively. The resulting lineage negative (Lin^-^) population was gated into three distinct subpopulations using rat-specific antibodies against the cell adhesion molecule CD24 (cluster of differentiation 24) and CD29 (integrin beta-1), confirming previous gating results ^28^ (Figure 3A, S3A). Three distinct populations were identified that represented luminal (CD24^hi^/CD29^mid^), basal (CD24^mid^/CD29^hi^), and stromal (CD24^low^/CD29^hi^) cells. We assessed the mammary subpopulations at PND 22 and 43 and observed that cell viability fluctuated by timepoint and genotype (Figure S3B). Due to the duration of time involved in MEC isolation, we suspected that cell viability may be affected and wanted to determine whether this influenced distinct cell populations or genotypes. To address this, cell-type population percent differences among the four genotypes, at each timepoint, were assessed via a beta mixed-effects model. To account for the effect of viability on the cell population percentage, which varied significantly as a function of cell-type, time point, and genotype (Figure S3C); a four-way interaction between all four variables was evaluated, but there was no evidence of a difference (p = 0.922). However, there was significant evidence for several three-way interactions: *viability x cell-type x genotype, timepoint x viability x cell-type,* as well as a *timepoint x cell-type x genotype* (Figure S3C; p = 0.008, p < 0.0001, and p < 0.0001, respectively). Therefore, the final model included these three three-way interactions, their composite two-way interactions, and individual main effects. This complex adjustment of viability properly accounts for any potential confounding effect cell-type viability may have had on the results. Using this model, we discovered there was a significant increase in luminal cells in the *Nf1^IF/+^* and *Nf1^IF;PS-21/+^* glands at 22 days, yet at 43 days, a significant decrease in the luminal population was detected in the *Nf1^PS-20/+^* and *Nf1^IF;PS-21/+^* glands (Figure 3B). For the basal population, a significant increase in the basal cell population was observed in *Nf1^IF;PS-21/+^* glands at 22 days, while a positive trend was present in *Nf1^IF/+^* and *Nf1^PS-20/+^* glands (Figure 3C). By 43 days, both the *Nf1^PS-20/+^* and *Nf1^IF;PS-21/+^* glands had a significant increase in basal population countered by a significant decrease in the luminal population. The results demonstrate a preferential expansion of the basal cell compartment between 22 and 43 days in *Nf1*-deficient glands. Notably, the *Nf1^PS-20/+^* and *Nf1^IF;PS-21/+^* rats that have a shift in luminal-basal lineages also have the most aggressive phenotype.^20^ We observed a decrease in the basal cell population at 43 days in *Nf1^IF/+^* mammary glands, which may produce the delayed tumor onset observed in this line (~10 mos). Collectively, these data reveal a shift in luminal to basal lineage commitment within the *Nf1*-deficient lines that have the earliest tumor onset.

**Figure 3:**
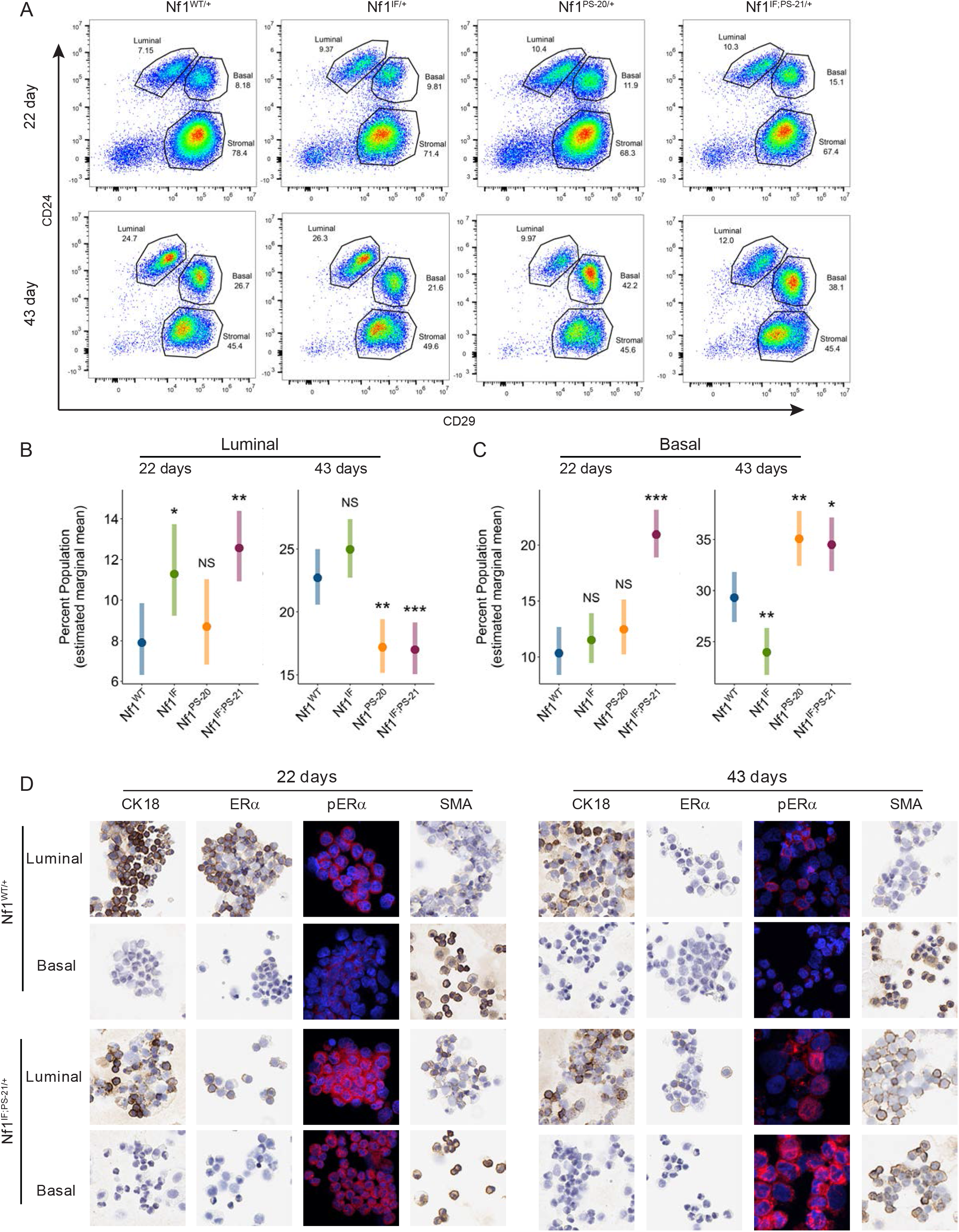
Mammary lineage population and marker expression changes in *Nf1*-deficient rats. A) Flow cytometry analysis of CD24 (luminal marker) and CD29 (basal marker) in digested mammary fat pads at 22 and 43 days in wild-type versus *Nf1*-deficient rats. B, C) 95% confidence intervals for mean percent population of flow cytometry sorted B) luminal cells and C) basal cells at 22 and 43 days per genotype (n=5 animals per genotype; *p < 0.05, **p < 0.01, ***p < 0.001). D) Immunostaining of luminal and basal cells spun onto glass slides following FACS of wild-type and *Nf1*-deficient female 4^th^ mammary fat pads. Cells were stained for CK18, ER, pER, and SMA. Images were taken at 200x (IHC) and 600x (IF) magnification and insets are shown.

To validate these results, we assessed lineage specific protein expression in each genotype at 22 and 43 days. Following FACS, cytospun cells were immunostained with epithelial lineage markers (Figure 3D). As expected, *Nf1^WT/+^* luminal cells were CK18+ and basal cells were SMA+ at both 22 and 43 days; whereas *Nf1^IF;PS-21/+^* luminal cells had decreased CK18 and mild SMA expression at both timepoints. We also assessed ER which is normally expressed in luminal populations. In addition, we examined phosphorylated S118 ER, the ERK phosphorylation site that facilitates coactivator interactions and can be impacted by RAS dysregulation. ^33,34^ *Nf1^WT/+^* luminal cells were ER+ and pER+ at 22 days, yet decreased ER+ and pER+ expression was observed at 43 days, whereas basal cells were negative for both ER and pER. In *Nf1^IF;PS-21/+^* glands pER (S118) was strongly expressed in both luminal and basal cells. Together, the shift in luminal to basal populations and the co-expression of luminal and basal lineage markers within *Nf1*-deficient mammary epithelium demonstrates that *Nf1* functionality impacts mammary lineage commitment.

### *Nf1* deficiency promotes lineage plasticity within mammary glands

The evidence that *Nf1* deficiency and the subsequent loss of neurofibromin function alters the mammary epithelial lineages in early development is illustrated in 1) the coexpression of luminal/basal markers in the TEB body (Figure 2C), the shift in luminal to basal populations (Figure 3B), and the altered SMA and ER expression in sorted luminal and basal populations (Figure 3D). To examine the impact of *Nf1* deficiency on cell fate later in mammary development and throughout tumorigenesis, we immunostained *Nf1*^*WT*/+^, *Nf1*^*PS-20*/+^, or *Nf1*^*IF;PS-21*/+^ mammary glands at several timepoints with E-cadherin and SMA (Figure 4). To identify cells that express both luminal and basal markers, we used an image calculator in FIJI. For E-cadherin and CK14 image masks, channel thresholding with predetermined threshold values (based on multiple training sets of images taken from all genotypes) was applied to identify areas of lineage marker overlap. In mammary glands from PND 22 females, we observed few dual positive (Ecad^+^/CK14^+^) cells within the *Nf1*^*PS-20*/+^ ducts; however, a substantial number of Ecad^+^/CK14^+^ cells were present in the *Nf1*^*IF;PS-21*/+^ ducts (Figure 4A). Next, we evaluated mammary glands at >50 days, in which the *Nf1*-deficient glands had hyperplastic changes but had not developed tumors (Figure 4B). At >50d, the *Nf1*^*PS-20*/+^ and *Nf1*^*IF;PS-21*/+^ glands exhibited varying levels of Ecad^+^/CK14^+^ cells, yet these dual-positive cells were not present in any *Nf1*^*WT*/+^ glands (Figure 4B). As opposed to the patterns observed at PND 22, the Ecad^+^/CK14^+^ cells in the >50d glands were consistently found at the periphery of the ductal structures corresponding with the basal cell layer.

**Figure 4:**
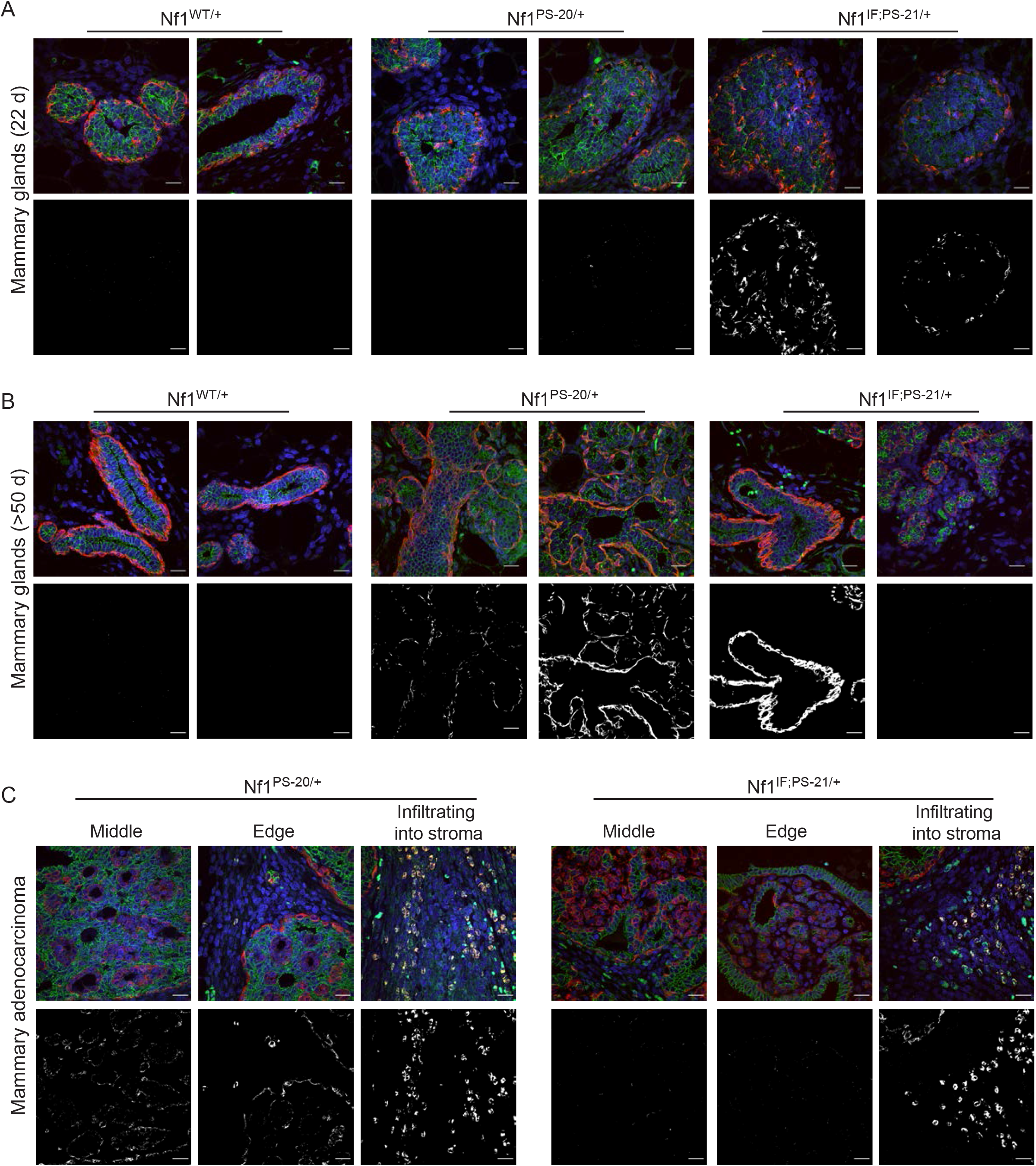
Dual lineage marker expression increases during development in *Nf1*-deficient mammary glands. Tissue sections of the 4^th^ mammary fat pad from wild-type and *Nf1*-deficient female rats were immunostained with CK14, E-cadherin, and DAPI at A) 22 days old, B) > 50 days old, and C) after mammary adenocarcinoma development. The top row in each panel represents merged CK14, E-cadherin, and DAPI staining. The bottom row in each panel represents FIJI image calculation of areas with CK14 AND E-cadherin overlay (white), indicating co-expression. Two replicates per genotype are shown. Images were acquired at 600x magnification, scale bar = 21 μm.

Next we examined whether these Ecad^+^/CK14^+^ cells persisted throughout tumor development. The expression patterns that we observed in the hyperplastic glands led us to question whether Ecad^+^/CK14^+^ cells are present within different tumor regions. To evaluate localization of the luminal/basal populations, we imaged the middle, edge, and invasive edge of each tumor. In an *Nf1^PS-20/+^* tumor, we observed few Ecad^+^/CK14^+^ cells in the middle of the tumor and edge of the tumor, but as in the hyperplastic glands, the Ecad^+^/CK14^+^ cells were located in either the basal layer or at the invasive edge of the tumor (Figure 4C, left panel). Strikingly, the strongest expression overlap was observed in tumor cells that had infiltrated into the stroma. This was also observed in an *Nf1^IF;PS-21/+^* tumor where Ecad^+^/CK14^+^ cells were identified in the stroma infiltrates, yet very weak signal was detected Ecad^+^/CK14^+^ cells within the tumor middle and invasive edge (Figure 4C, right panel). We then examined these luminal/basal subpopulations within one tissue section that contained a progression of normal, hyperplastic, and multiple adenocarcinoma region (Figure S4). As before we observed Ecad^+^/CK14^+^ cells in the hyperplastic gland and some tumor regions, but interestingly Ecad^+^/CK14^+^cells were not observed in tumors 4 and 5. Together these results demonstrate that Ecad^+^/CK14^+^ cells are present in *Nf1*-deficient mammary glands at the earliest stages of development, yet they are a minute subpopulation. In the earliest stages of tumor development, these luminal/basal populations are located at the basal layer, yet later are only observed at the invasive edge and infiltrating cells. These findings imply that *Nf1*-deficient tumors contain a subpopulation of cells with lineage plasticity between the luminal-basal epithelial states.

### *Nf1* mutation alters neurofibromin expression and localization throughout mammary gland and tumor development

As presented previously^20^, each of our *Nf1*-deficient lines correspond to variable levels of neurofibromin loss of function that correlate to tumor latency. To understand how these mutations alter neurofibromin function within the mammary gland, we performed confocal imaging of neurofibromin using an N-terminal antibody (Figure 5A). Within 21-day old *Nf1^WT/+^*glands, neurofibromin expression was localized to the TEB periphery and colocalized with SMA expression (Figure 5B). This expression pattern dramatically changed in the *Nf1*-deficient glands in which neurofibromin expression was present throughout the TEB body compartment. Even though neurofibromin/SMA colocalization was still observed at the periphery and within SMA+ body cells, diffuse neurofibromin expression was observed throughout the TEB. Closer examination of subcellular localization in *Nf1^WT/+^* glands (Figure 5B, insets) revealed that neurofibromin expression is primarily observed at the plasma membrane and weakly cytoplasmic. Neurofibromin membrane localization was completely distorted in *Nf1^IF/+^* and *Nf1^IF;PS-21/+^* tumor glands but was present in some cells within the *Nf1^PS-21/+^* TEBs.

**Figure 5:**
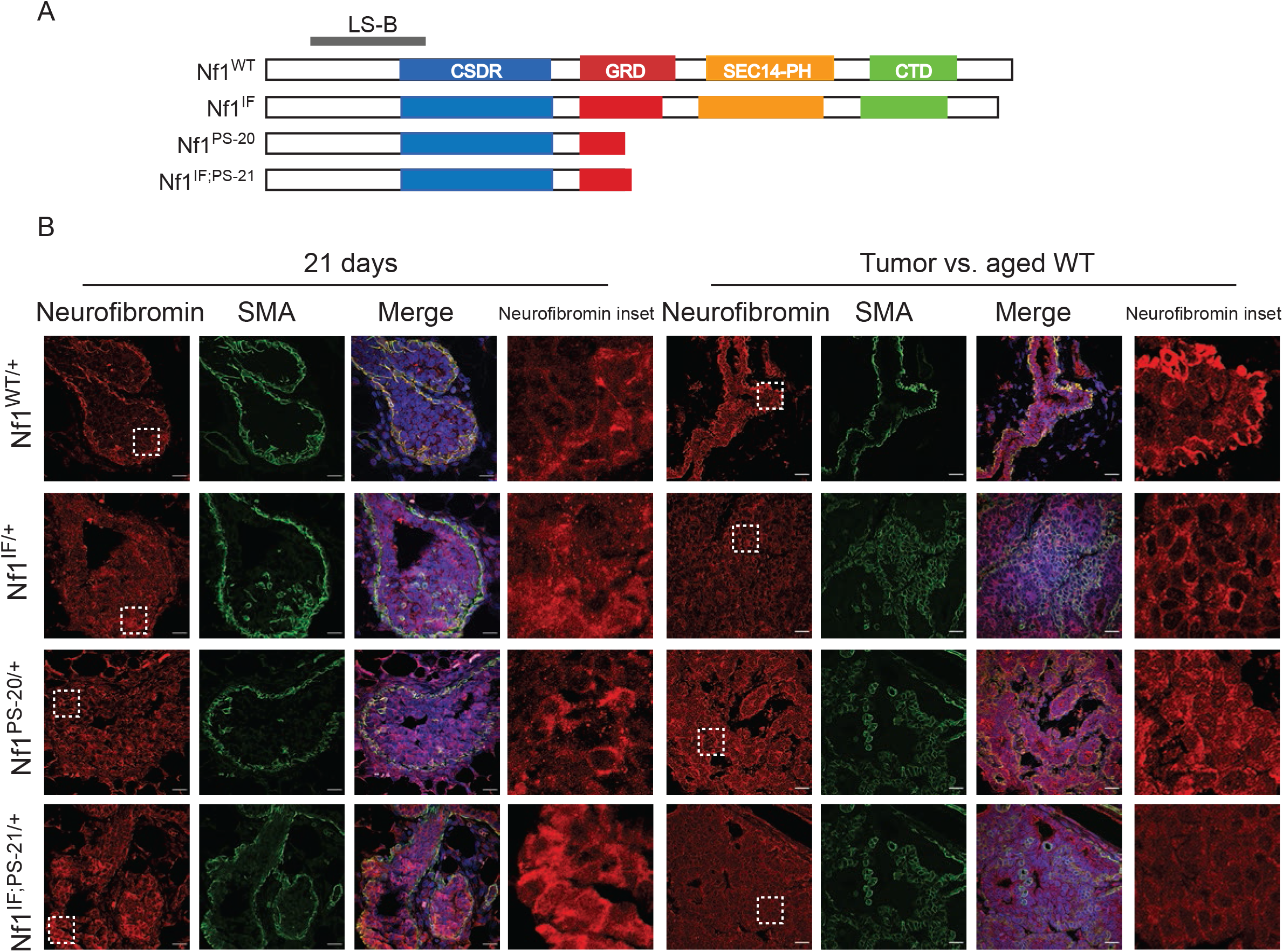
Neurofibromin expression in the mammary gland and breast tumors. A)Schematic of the LS-B antibody epitope in neurofibromin. B) Immunostaining of neurofibromin using the LS-B antibody in wild-type and *Nf1*-deficient mammary TEBs (21 days) and ducts/tumor in aged females from the 4^th^ mammary fat pad. Images were acquired at 600x magnification, scale bar = 21 μm.

In the aged *Nf1^WT/+^* glands, we observed intense membrane localization of neurofibromin at the ductal edge and weak cytoplasmic expression within the luminal layer (Figure 5B inset). In *Nf1-*deficient tumors, neurofibromin expression was present throughout the tumor and small neurofibromin^high^ subpopulations overlapped with high SMA expression. Membrane localization of neurofibromin was highest in *Nf1^IF/+^* tumors, yet both cytoplasmic and membrane expression was observed in *Nf1^PS-20/+^* and *Nf1^IF;PS-21/+^* tumor glands. Collectively, these results demonstrate that *Nf1* mutation alters expression and subcellular localization of neurofibromin in early mammary development which is present throughout tumorigenesis.

### *Nf1* deficiency restricts luminal progenitor potential *in vitro*

To assess the functional impact of *Nf1* deficiency on clonogenicity and mammosphere development/differentiation, we utilized three distinct *in vitro* assays, the colony forming cell (CFC) assay, the mammosphere assay, and the 3D growth assay. These assays are commonly used to measure stem and progenitor cell potential in human and mouse mammary epithelial cells ^35–37^, yet to our knowledge this is the first time they have been utilized for rat mammary cells. In the CFC assay, morphology and lineage marker expression in these colonies can be scored as basal-restricted, luminal-restricted, or mixed colonies that likely originate from basal, luminal, and bi-potent progenitors, respectively.^38^ We flow-sorted luminal or basal cells from 43-day old wild-type and *Nf1*-deficient female rats and immunostained with CK14 (red), E-cadherin (green), and DAPI (blue) (Figure 6A). In the luminal-sorted colonies, luminal-restricted (Ecad+) and mixed (Ecad+/CK14+) colonies were present in the *Nf1^WT/+^* cells as expected (Figure 6B, upper panel). We observed a decrease in mixed colonies within the luminal-sorted *Nf1^IF/+^* and *Nf1^PS-21/+^* populations, yet no mixed colonies in *Nf1^IF;PS-21/+^* populations were seen. In the *Nf1^IF;PS-21/+^* line, we also observed a decrease in luminal-restricted colonies. In the basal-sorted colonies, there was no substantial difference observed in the basal-restricted (CK14+) or mixed colonies between the *Nf1^WT/+^, Nf1^PS-20/+^* and the *Nf1^IF;PS-21/+^* lines (Figure 6B, lower panel). However, we observed a decrease in luminal-restricted colonies in the *Nf1^PS-20/+^* and *Nf1^IF;PS-21/+^* lines and in contrast an increase in luminal-restricted colonies in the *Nf1^IF/+^* line. There were no mixed colonies in the basal-sorted *Nf1^IF/+^* line. This luminal differentiation pattern mirrors what we observed in the flow-sorted luminal populations at 43 days with an increase in luminal populations in the *Nf1^IF/+^* line and a decrease in the *Nf1^PS-20/+^* and *Nf1^IF;PS-21/+^* lines (Figure 3B).

**Figure 6:**
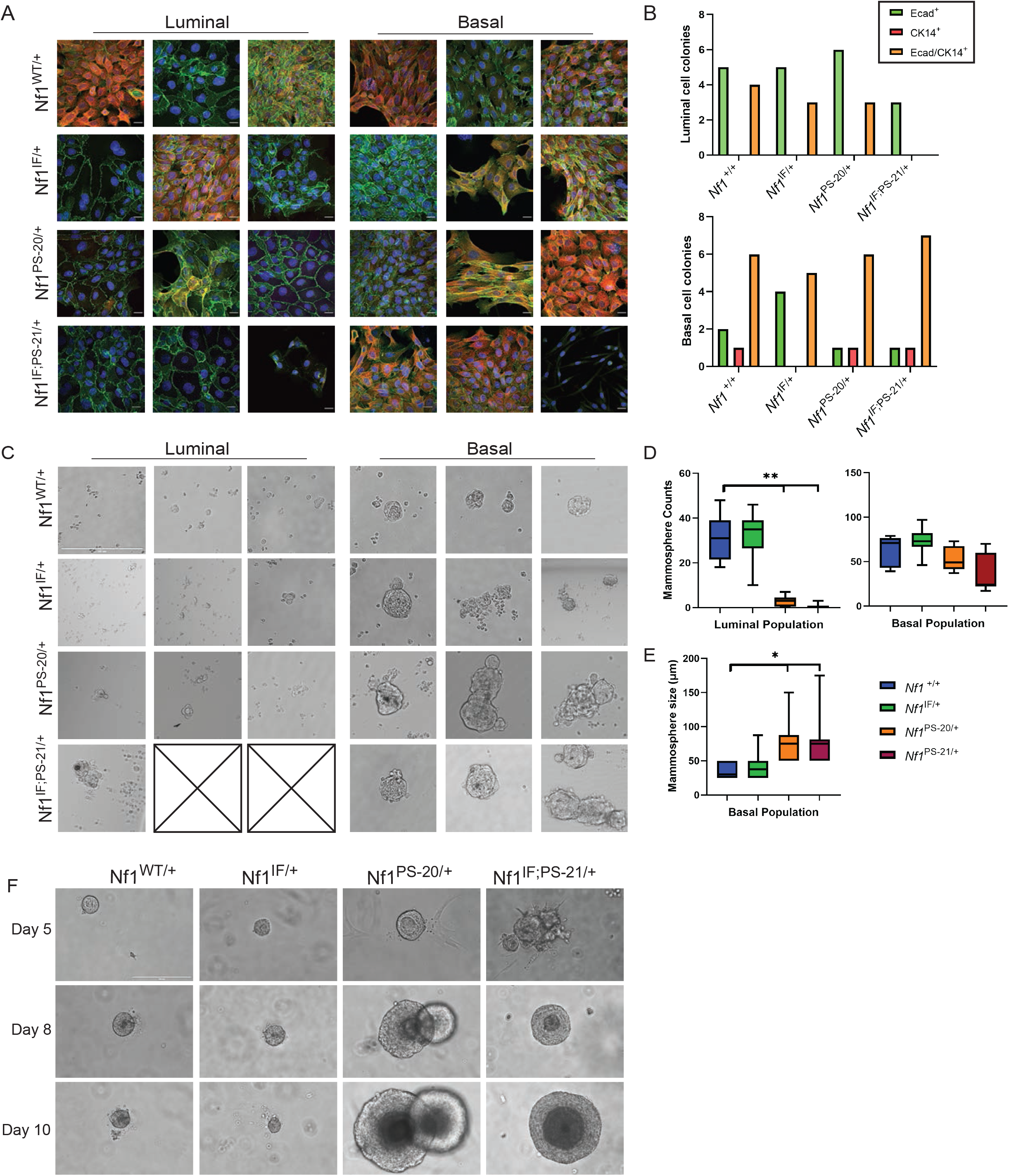
*Nf1*-deficient stem and progenitor cells display altered functionality. A)Immunostaining with CK14, E-cadherin, and DAPI of colonies grown from flow sorted luminal or basal cells. Images were acquired at 600x magnification, scale bar = 21 μm. B) Luminal and basal colony counts based on E-cadherin^+^ (green), CK14^+^ (red) or Ecad/CK14^+^ (orange) lineage marker expression. C) Representative images of mammospheres formed from 20,000 flow sorted cells per population in ultra-low attachment plates. Images were taken at 200x magnification, scale bar = 200 μm. Boxes with an x indicate samples where none of the replicates formed mammospheres. D) Quantitation of mammospheres formed from 180,000 flow sorted cells per population (60,000 cells/population/animal; n = 3; **p < 0.0001). E) Quantitation of mammosphere size from flow sorted mammary basal cells. 25 spheres per genotype were measured from images taken at 100x magnification (*p < 0.05). F) Representative images of MECs, from digested 4^th^ mammary fat pads of wild-type and *Nf1*-deficient female rats, grown for 5, 8, 10 days in Matrigel. Images were taken at 200x magnification, scale bar = 200 μm.

We used mammosphere and 3D-colony culture assays to evaluate mammary stem/progenitor potential based on their ability to form mammospheres in ultra-low attachment conditions.^35,39^ All luminal mammospheres were small acinar-like structures, which is indicative of mature and progenitor luminal cells (Figure 6C).^40^ Mammosphere formation was significantly decreased in the luminal-sorted *Nf1^PS-20/+^* and *Nf1^IF;PS-21/+^* cells (Figure 6D, p < 0.0005). In 3D-colony assays, basal/stem cells are known to form large, disorganized spheres that exhibit lateral budding or branching outgrowths as we observed in *Nf1^WT/+^* cultures.^40^ Mammospheres from the *Nf1*-deficient lines were larger than *Nf1^WT/+^* mammospheres, and they were commonly multi-lobed, indicating hyperproliferation. Although there was no significant difference in the number of basal-sorted *Nf1*-deficient mammospheres, the size of the mammospheres generated from *Nf1^PS-20/+^* and *Nf1^IF;PS-21/+^* were significantly larger than those from *Nf1^WT/+^* (Figure 6D-E, p < 0.05). Lastly, we isolated mammary epithelial cells and cultured them in Matrigel, where MaSCs have the ability to form solid, pleomorphic structures, and luminal progenitors form hollow acinar structures ^36,41–45^. Surprisingly, very few of the luminal 3D colonies, even in the *Nf1^WT/+^,* had hollow lumens but most were small and translucent indicating alveolar-type structures (Figure S5). We also observed that many of the mammospheres from the *Nf1^PS-20/+^* and *Nf1^IF;PS-21/+^*luminal cells were unable to form any 3D colonies. The basal 3D colonies from all genotypes were a mix of optically dense, large, and complex structures with a plethora of invasive processes/protrusions and simple solid colonies (Figures 6F and S5).

### *NF1* deficiency transforms human breast cells

To evaluate the effect of *NF1* loss of function in human breast cells, we used the immortalized, nontumorigenic human breast epithelial cell line MCF10A. CRISPR guides targeting exon 21 within the GRD domain were constructed and used either as single or dual-guide RNAs with Cas9 nuclease (Figure S6A). We established three cell lines (i.e., g1, g2, g1/2) and *NF1* loss was validated by PCR and Western blot analysis (Figure S6B). MCF10A cells were grown in 3D culture conditions and as expected, the empty vector (EV) MCF10A line developed small spherical 3D colonies (Figure 7A). In contrast, MCF10A lines with *NF1* mutations had distinct phenotypes, with the NF1-g2 and NF1-g1/g2 lines developing larger, grape-like clusters.

**Figure 7:**
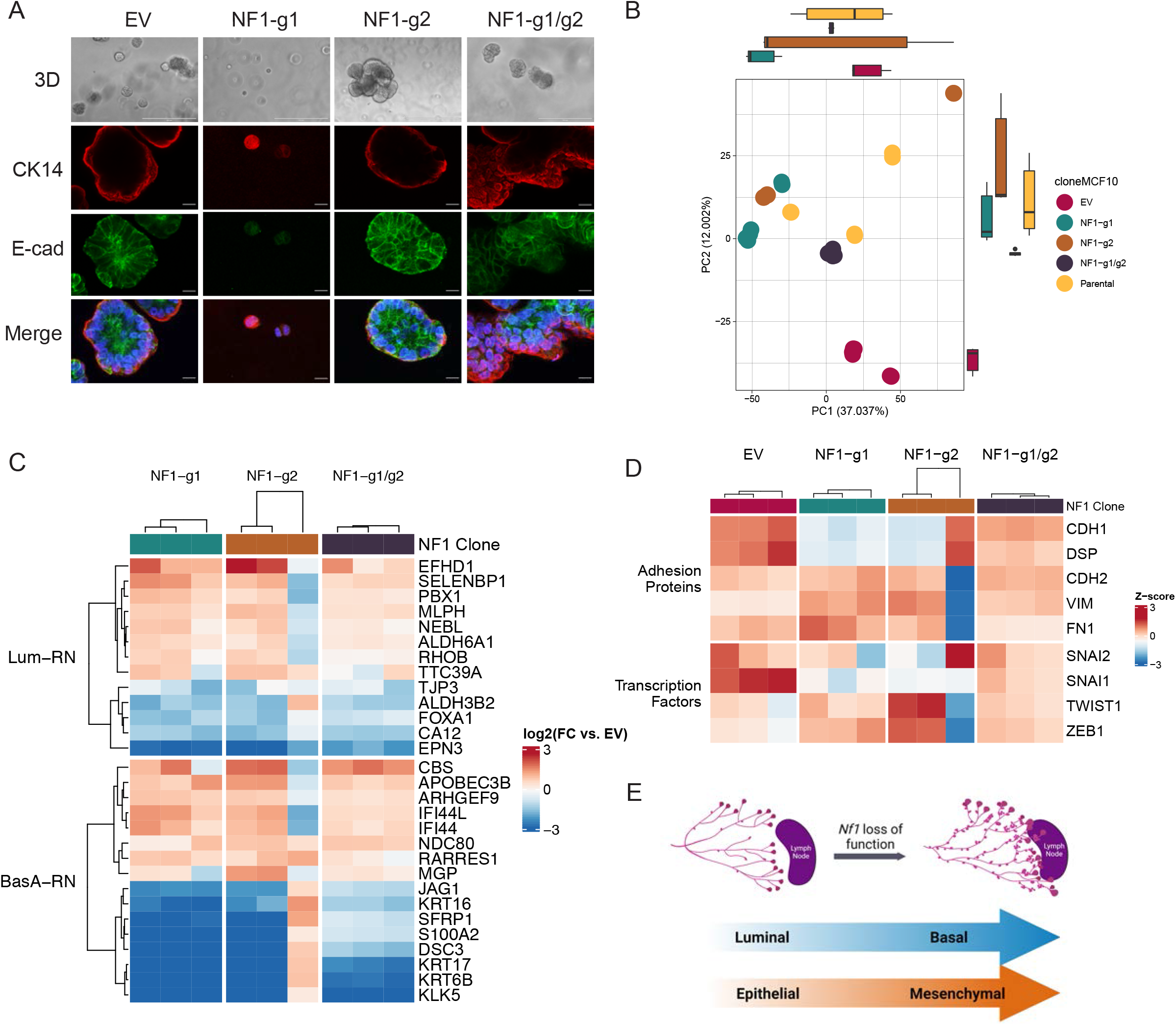
*NF1* deficiency induces EMT. A) MCF10A *NF1* CRISPR mutants and EV control were grown in Matrigel for 10 days and then stained with CK14, E-cadherin, and DAPI. Images were taken at 600x, scale bar = 21μm. B) principal component analysis controls showing variation in samples from RNA sequencing of MCF10A *NF1* CRISPR mutants compared to empty vector and parental controls. C&D) Heatmap of *NF1* CRISPR mutants (log fold change vs. the empty vector control) showing C) consistent changes in RNA expression in both luminal and basal gene signatures and D) changes in RNA expression of proteins related to EMT. E) Schematic showing *Nf1* loss of function leads to accelerated mammary ductal outgrowth, lineage plasticity between luminal and basal cells, and pushes cells toward a mesenchymal state.

Interestingly, the NF1-g1 line formed few, smaller 3D colonies with the majority surviving as single cells. Immunostaining showed changes in expression of E-cadherin and CK14 in the NF1-g2 and NF1-g1/g2 cells. NF1-g1 colonies weakly expressed E-cadherin and strongly expressed CK14 suggesting these cells have become more basal-like. We performed RNAseq and utilized principal component analysis (PCA) to visualize the variability across the *MCF10A-NF1^mut^* cell lines (Figure 7B). PCA displayed the clonal diversity in the parental MCF10A lines and the global shift in gene expression in the *NF1*-mutant lines compared to the EV. Next we examined the impact of *NF1* deficiency on the MCF10A transcriptomes within previously defined luminal and basal classifier gene signatures ^46,47^ and observed that *NF1* deficiency induced consistent expression changes in both luminal and basal gene sets (Figure 7C). MCF10A-*NF1^mut^* lines upregulated several genes in the luminal signature known to promote cancer progression or correlate with poor prognosis (*EFDH1, MLPH, ALDH6A1*)^48–50^ Notably, several genes were downregulated that are involved in ERα signaling and cofactor regulation (*PBX1, RHOB, FOXA1, CA12).* In the basal gene signature, MCF10A-*NF1^mut^* lines showed downregulation of genes involved in cell adhesion (*KRT6B, KRT17, DSC3, JAG*). Two of the most upregulated genes in the basal signature are metabolic genes that protect against oxidative stress (*CBS*)^51^ and are involved in the pyrimidine salvage pathway (*AOBECD3B*).^52,53^ We observed similar gene expression changes in the MCF10A-*NF1^mut^* lines except in one subclone in the NF1-g2 line (Figure 7B-C). Because of the Ecad^+^/CK14^+^ cells we observed in *Nf1*-deficient tumors (Figure 4C), we evaluated the expression changes in several genes that are essential for EMT^54,55^ and saw that the clones that often did not form 3D mammospheres (NF1-g1 and NF1-g2) had less *CDH1* (E-cadherin) and *DSP* (Desmoplasia) expression but had increased mesenchymal adhesion protein (*CDH2, VIM,* and *FN1*) and EMT transcription factor (*TWIST1* and *ZEB1*) expression (Figure 7D). Many of the genes that were deregulated in the luminal, basal, and EMT gene sets were among the most differentially expressed genes in MCF10A-*NF1^mut^* lines (Figure S6C). Using gene set enrichment analysis (GSEA), we found that several KEGG pathways were significantly enriched in the *MCF10A-NF1^mut^* lines compared to the EV control, such as pathways in cancer (includes RAS signaling), regulation of actin cytoskeleton, focal adhesion, and cytokine-cytokine receptor interaction (Figure S6D). The enrichment of these pathways in *NF1*-deficient cells aligns with the EMT signatures (Figure 7D) and gene expression changes we observed (Figure 7C).

## DISCUSSION

Over the last ten years, both genomic and epidemiological studies have uncovered *NF1* as a key tumor suppressor in both sporadic and *NF1*-related breast cancer. Because of the prevalence of neuronal tumors in NF patients, much of our understanding of the functional loss of *NF1* is derived from tumorigenesis studies within nerve and brain tissues. The recent studies demonstrating a role for *NF1* in endocrine-resistance^3,4,6^, the high incidence of breast cancer in NF individuals^10,56^, and loss of *NF1* in resistance to RAS pathway inhibition in other cancers^57,58^, highlight the fundamental need for a more comprehensive understanding of *NF1* functions within the breast and other tissues. To interrogate the role of *NF1* in mammary homeostasis and tumor initiation, we utilized our immunocompetent rat model of *Nf1* deficiency. The distinct *Nf1* indels permitted us to explore how partial and severe loss of neurofibromin function impacts mammary morphogenesis and homeostasis. We demonstrate that *Nf1* deficiency dramatically accelerates mammary morphogenesis and alters TEB cell organization and proliferation. Notably, the most considerable mammary gland alterations were observed in the *Nf1*-deficient lines with the most aggressive tumor phenotypes, *Nf1^PS-20/+^* and *Nf1^IF;PS-21/+^* lines.^20^ These striking results phenocopy the accelerated mammary development caused by haploinsufficiency of the ER co-repressor REA^59,60^ which substantiate the recent discovery that neurofibromin acts as an ER co-repressor.^61^

In addition to accelerated development and altered TEB organization, we identified a shift in luminal to basal epithelial lineage commitment within the *Nf1*-deficient lines that have the earliest tumor onset. Lineage plasticity was first detected within the *Nf1^PS-20/+^* and *Nf1^IF;PS-21/+^* TEBs where diffuse E-cadherin expression was observed throughout the TEB in addition to dual positive SMA+/Ecad+ cells within the TEB body. FACS analysis demonstrated that in later stages of mammary development there was a significant increase in the basal epithelial population corresponding to a decrease in the luminal population. In contrast, the *Nf1^IF/+^* mammary glands, which have a delayed tumor onset (~10 mos) compared to *Nf1^PS-20/+^* and *Nf1^IF;PS-21/+^* rats, demonstrated a significant decrease in the basal population. In addition, we observed altered expression of pER (S118) with a shift in luminal to basal populations in 43-day *Nf1*-deficient glands. Together these results allude to an inherent lineage plasticity in the aggressive *Nf1^PS-20/+^* and *Nf1^IF;PS-21/+^* lines compared to the delayed *Nf1^IF/+^* breast cancer model. We hypothesize that lineage plasticity corresponds to level of neurofibromin function. These results emphasize the need for a more thorough understanding of neurofibromin’s functional domains beyond the RAS regulatory domain.

Several foundational studies have revealed that cell plasticity is often a critical player in tumor initiation. Early studies identified specific progenitor cells as the cell of origin for breast cancer subtypes; however, recent work has demonstrated that intrinsic plasticity is important for our understanding of both tumor origins and heterogeneity.^54^ For example, a genetic approach targeting *Pik3ca^H1047R^* to basal (K5CreER) or luminal (K8CreER) populations demonstrated that expression of *Pik3ca^H1047R^* within basal cells resulted in ER^+^/PR^+^ luminal tumors, whereas expression of *Pik3ca^H1047R^* within luminal cells led to basal ER^-^/PR^-^ tumors.^62^ Similarly, studies in *Brca1*-mutated mouse models and human tissues have demonstrated that *Brca1* mutation can abrogate luminal progenitor differentiation resulting in basal breast cancer. These studies reveal the inherent lineage plasticity within the mammary epithelium that leads to cancer initiation.^43^ Our findings demonstrate that *Nf1* loss of function promotes luminal-basal plasticity within the developing mammary gland (Figure 7E). The major challenge in delineating the tumor cell of origin further in these models is the lack of validated rat antibodies for distinguishing luminal and basal progenitor populations. As CRISPR-edited rat models become more prevalent due to their human-like pharmacokinetic profiles and mammary development^63–65^, it is likely that rat-specific resources will be rapidly developed that will allow more extensive analysis of lineage populations. In this study, to clarify the impact of *Nf1* function on lineage commitment and progenitor capacity, we utilized several *in vitro* assays. The most striking observation was the decreased mammosphere formation in luminal-sorted *Nf1^PS-20/+^* and *Nf1^IF;PS-21/+^* cells and the coinciding increase in mammosphere size in basal-sorted *Nf1*-deficient cells. Collectively, our in vitro and in vivo results suggest that *Nf1*-deficiency restricts luminal progenitor potential and a shift towards the basal lineage.

EMT is well characterized example of phenotypic plasticity and has been established to be involved in various stages of tumor progression and metastasis in diverse cancer types.^54^ Recently, several studies have demonstrated that tumor cell populations can exist within hybrid EMT stages that provide different levels of cell plasticity to enhance metastasis or treatment resistance.^66^ Hybrid EMT states often co-express epithelial and mesenchymal markers, such as cells within invasive breast cancers that co-express E-cadherin and vimentin.^67^ Similarly in our analysis of *Nf1*-deficient tumor progression, we observed Ecad^+^/CK14^+^ co-expression in subpopulations within both hyperplastic and neoplastic tissues. In the earliest stages of tumor development, these luminal/basal populations were located at the basal layer, yet later Ecad^+^/CK14^+^ cells were only observed at the invasive edge and stromal-infiltrating cells. These findings imply that *Nf1*-deficient tumors contain a hybrid EMT subpopulation in which this phenotype plasticity drives invasiveness.

Together our findings support a model in which *NF1* loss of function results in lineage plasticity during the early stages of mammary morphogenesis (Figure 7E). This study reveals a previously unknown role for the tumor suppressor *NF1* in mammary homeostasis and phenotypic plasticity during tumor progression and may indicate the driver behind high breast cancer risk and poor outcomes in individuals with NF.

## STAR Methods

Detailed methods are provided in the online version of this paper and include the following:

### KEY RESOURCES TABLE

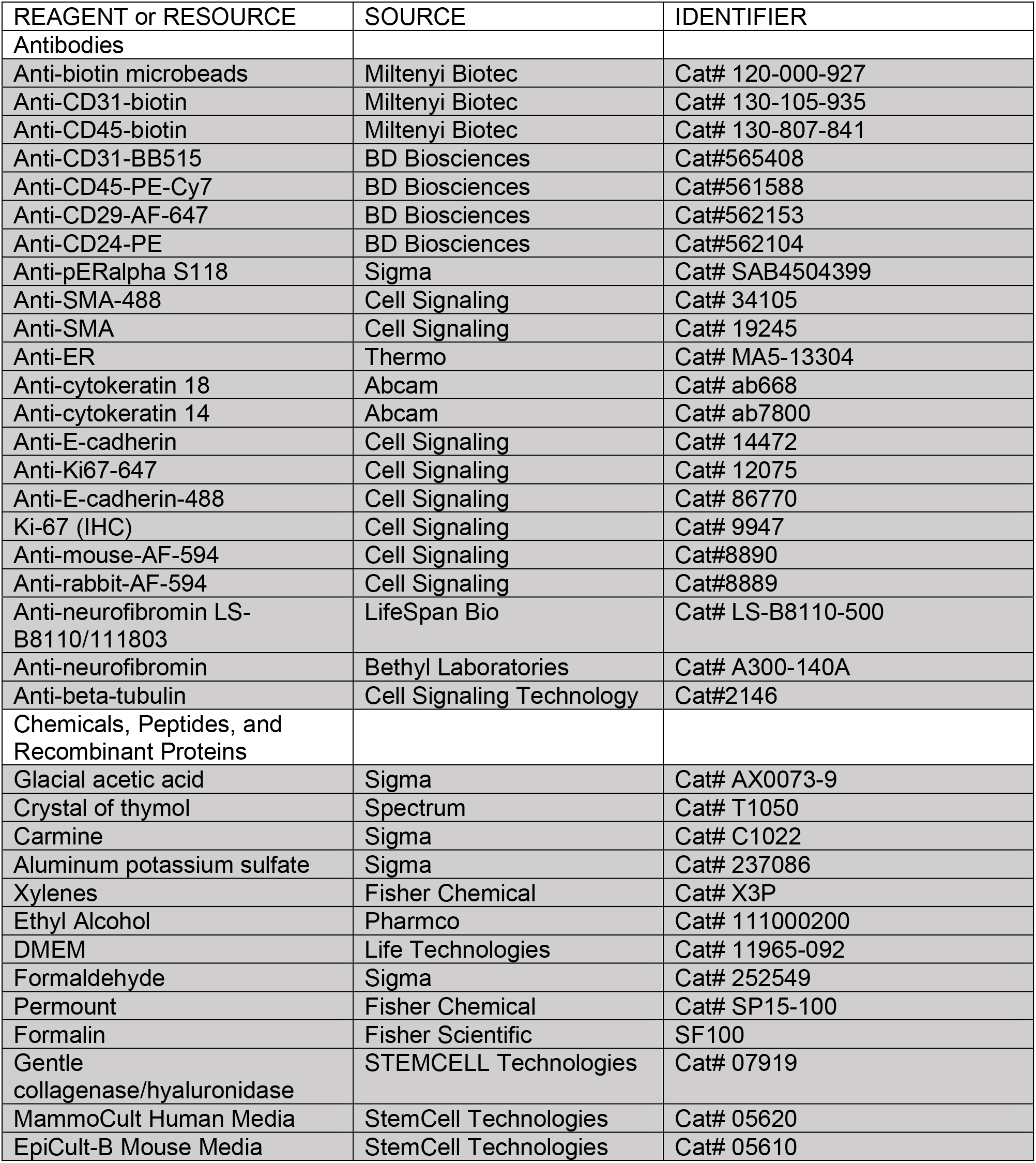
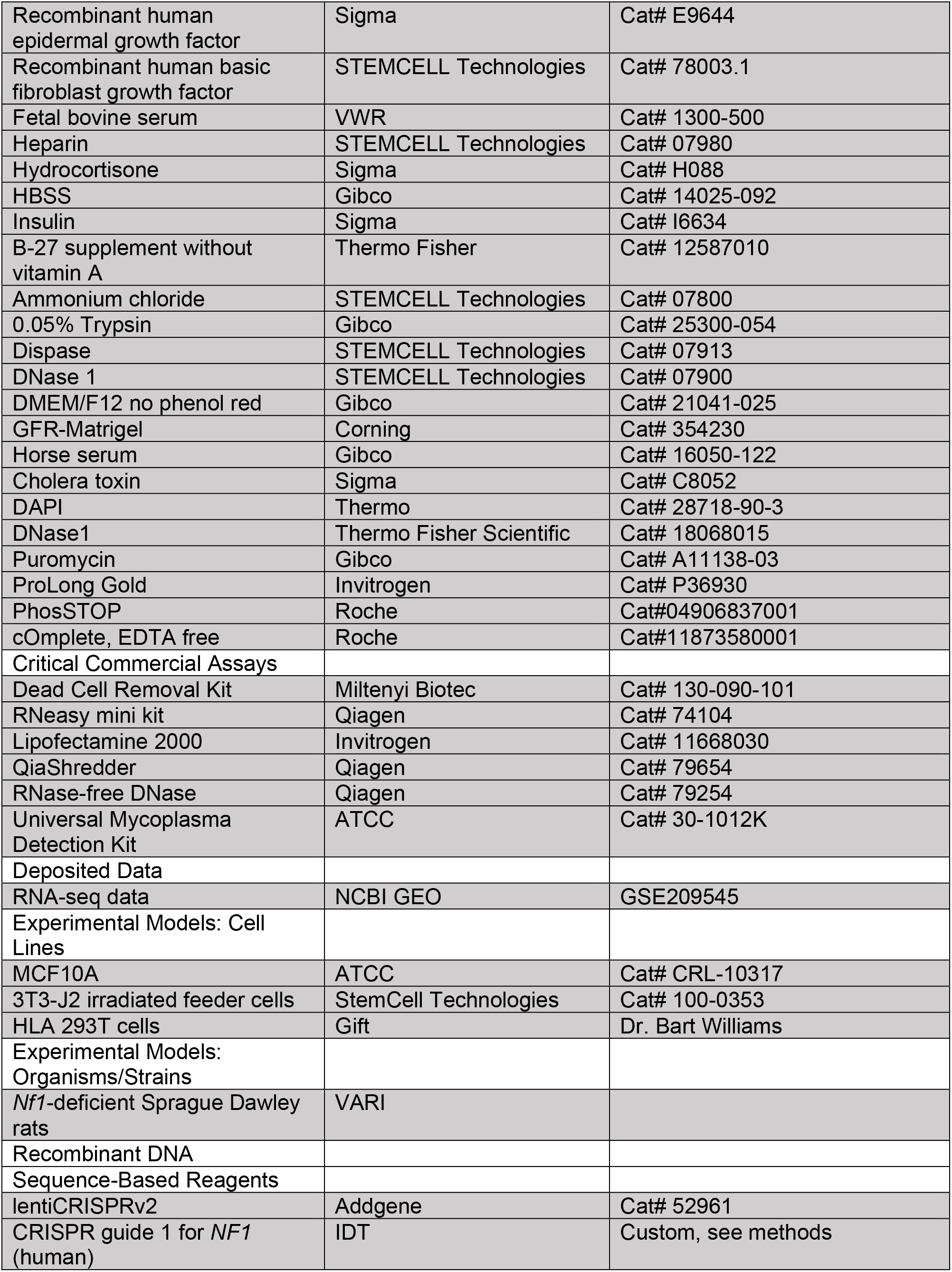
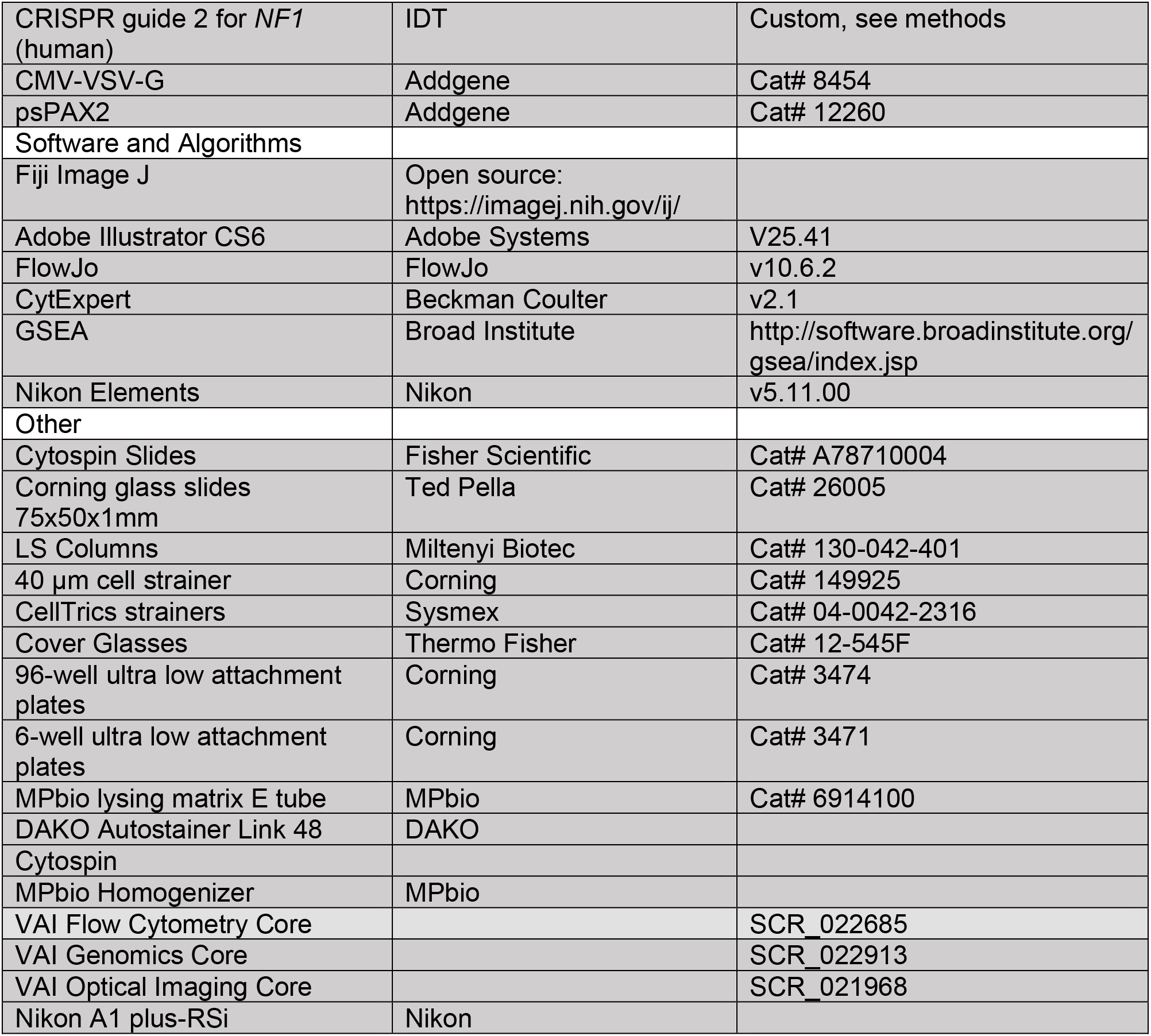

### CONTACT FOR REAGENTS AND RESOURCE SHARING

Further information and requests for resources and reagents should be directed to and will be fulfilled by the Lead Contact, Dr. Matt Steensma.

### MATERIALS AVAILABILITY

Plasmids generated in this study have been deposited to Addgene:

NF1-lentiCRISPRv2-g1

NF1-lentiCRIPSRv2-g2

### EXPERIMENTAL MODEL AND SUBJECT DETAILS

#### Animals

Female and male immunocompetent *Nf1*-deficient Sprague Dawley rats^20^ were bred in house at the Van Andel Research Institute. Genotypes include *Nf1^+/+^*; *Nf1^PS-20/+^*, *Nf1^IF/+^*, and *Nf1^IF;PS-21/+^*. Male *Nf1*-deficient rats were bred to wild-type female rats and female pups were used at the indicated developmental time-point. Animals were housed 2 to a cage, had *ad libitum* access to food and water, and were kept under a 12h light-12hr dark cycle. All animal protocols were approved by the Van Andel Research Institute Animal Care and Use Committee.

#### Cell Culture

Rat mammary epithelial cells derived from female 4^th^ mammary fat pads were grown in MEC media (phenol red-free DMEM/F12, 10% horse serum, 20ng/mL EGF, 0.5ug/mL hydrocortisone, 100ng/mL cholera toxin, and 10ug/mL insulin). Human MCF10A breast cells, derived from a female patient in 1984, were also grown in MEC media. All cells were grown at 37°C in a 5% CO_2_ incubator. Mycoplasma testing (ATCC) every 6 months confirmed cells were not infected.

### METHODS DETAILS

#### Carmine Whole Mounts

Following euthanasia, the fourth mammary fat pads of wild-type and *Nf1-*deficient rats were dissected from the skin and spread onto glass slides. The right fourth mammary fat pad (R4) was fixed in formalin for 48 hours, while the other pad (L4) underwent carmine staining. Glands were allowed to attach to the slides for 30 minutes at room temperature (RT) and then transferred, on the slide, to Carnoy’s fixative (75% glacial acetic acid and 25% absolute ethanol). Slides underwent one wash in 70% ethanol (1 hr., RT) and one wash with distilled water (30 min, RT). Slides were immediately placed in carmine alum stain (1g carmine; 2.5g aluminum potassium sulfate; dissolved in 500mL distilled water plus a crystal of thymol) for 2 days and then washed in increasing concentrations of ethanol (1 hr. each, 70% > 95% > 100%). Lastly, slides were delipidated in xylenes for at least two days. Whole mounts were imaged using a stereo microscope.

#### Mammary Fat Pad Digestion

Freshly isolated fourth mammary fat pads (MFP) were digested into a single cell suspension as described in Dischinger *et al.* (2018) and Tovar *et al.* (2020). Briefly, finely minced MFP was digested overnight at 37°C in EpiCult B media containing 5% FBS, 1% gentle collagenase/hyaluronidase, 10ng/mL recombinant human epidermal growth factor, 10ng/mL recombinant human basic fibroblast growth factor, and 4ug/mL heparin. Pelleted cells were incubated with ammonium chloride in HBSS + 2% FBS to lyse red blood cells followed by 0.05% trypsin, and then 5U/mL dispase + 0.1mg/mL DNase I to further digest cell clumps. Cells were then strained through a 40 μm cell strainer.

#### Isolation of Rat Mammary Epithelial Cells

Live rat mammary cells derived from a mammary fat pad digestion were separated from dead cells using Miltenyi’s Dead Cell Removal Kit. Then CD31-biotin and CD45-biotin antibodies were used to label and select out endothelial and hematopoietic cells using anti-biotin microbeads and a MACS separator magnetic with LS columns (Miltenyi). To deplete fibroblasts, cells were plated in ultra-low attachment for 48 hours in MEC media. After 48 hours, cells were plated on tissue-culture treated plates in MEC media with no cholera toxin.

#### Flow Cytometry Analysis and Sorting of Mammary Epithelial Cells

For flow analysis and sorting, 1×10^6^ digested mammary cells in 100 uL were stained for 30 minutes room temperature using the following antibodies: CD31-BB515 (BD, 0.125ug), CD45-PE-Cy7 (BD, 0.0625ug), CD29-AF-647 (BD, 0.03125ug) and CD24-PE (BD, 0.125ug). Cells were washed and then stained with DAPI and analyzed on a CytoFlex S flow cytometer or sorted on a MoFLO Astrios. Sorted cells were spun onto ringed glass slides using a Cytospin at a concentration of 1×10^5^ cells/slide.

#### Immunostaining

Rats were euthanized at the indicated time points and their fourth mammary fat pads harvested and fixed in 10% formalin for 48 hours for immunohistochemistry (IHC) or immunofluorescence (IF). Cells on coverslips were fixed and stained as previously described.^32^ Cells on cytospin slides were fixed in ethanol for 10 minutes and then stained by IHC or IF. High pH antigen retrieval and staining was performed on the DAKO Autostainer Link 48 on 5-micron tissue sections. All IF slides were stained with DAPI. Cells in Matrigel were fixed with formaldehyde, blocked, and targets were stained sequentially. DAPI was used to stain cell nuclei. Slides were mounted with ProLong Gold.

#### Confocal Microscopy

For imaging, at least three regions of interest (ROI) were imaged per sample using a 60x 1.4 N.A. Plan Apo VC oil immersion objective on a Nikon A1 plus-RSi laser scanning confocal microscope on an eclipse Ti base, with Nikon Elements acquisition software. The microscope is outfitted with DU4 high sensitivity detectors, solid state excitation lasers (405, 488, 561, and 640nm), and single bandpass emission filters (450/50, 525/50, 595/50, and 700/75nm). To image TEBs, structures were identified at 10X magnification (Nikon, 0.3 N.A., plan fluor, air immersion) before switching to the Nikon oil immersion 60x Plan Apo VC with 1.4 NA. Images were collected with a pixel density of 1024×1024, and a 0.5μm z-stack depth set to cover the tissue thickness. Laser power and PMT gain settings were optimized using control samples and held consistent for each data set. Maximum intensity projection TIFF images were generated using FIJI.

For E-cadherin and CK14 image masks, channel thresholding with predetermined threshold values based on multiple training sets of images taken from all genotypes was applied to generate areas of marker overlap with the image calculator function in FIJI. To avoid staining artifacts, we selected at least two tissue blocks from animals of each genotype and imaged at least three ROI per section. For tumors, the middle, edge, and invasive edge were imaged. In the case where E-cadherin and CK14 colocalization varied drastically among samples, we selected additional slides from another animal of the same genotype for comparison.

#### Western Blot

Cells were washed once with ice-cold PBS, scraped, and lysed in Rb buffer (20 mM TrisHCL, pH 7.6; 5 mM EDTA; 150 mM NaCl; 0.5% NP-40; 50 mM NaF; 1 mM beta-glycerophosphate) supplemented with PhosSTOP and protease inhibitors. Samples were resolved by SDS-PAGE and immunoblotting was carried out using antibodies against neurofibromin (Bethyl) and betatubulin.

#### Colony Forming Cell Assay

To identify progenitor cells that can form *in vitro* colonies we followed the methods in.^68^ Briefly we plated 1,250 flow sorted mammary cells into one well of a 24-well plate that had already been seeded with 2×10^4^ 3T3-J2 irradiated feeder cells (100-0353, StemCell Technologies) in MMS media supplemented with 5% FBS and 100ng/mL cholera toxin (Sigma). MMS media consisted of human MammoCult Complete Media (StemCell Technologies), 2% B27 without vitamin A (Invitrogen), 20ng/mL bFGF (Stem Cell Technology), 20ng/mL EGF (Sigma), 10μg/mL Heparin (StemCell Technologies), 10μg/mL Insulin (Sigma), and 1μg/mL hydrocortisone (Sigma). 24 hours after seeding, cells were switched to serum-free MMS media and left to form colonies for 9 days. Three wells were plated for both the luminal and basal flow sorted populations for each animal (3 animals per genotype). At the end of 9 days, colonies were processed for immunostaining. The number of colonies positive for E-cadherin, CK14, or both were counted for both the luminal and basal populations.

#### Mammosphere Assay

To identify mammary stem/progenitor cells, we utilized the mammosphere assay which has been described previously. ^35,68^ Briefly, 20,000 flow cytometry sorted basal or luminal cells were cultured for 9 days in MMS media in 96-well ultra-low attachment plates. MMS media consisted of human MammoCult Complete Media (StemCell Technologies), 2% B27 without vitamin A (Invitrogen), 20ng/mL bFGF (Stem Cell Technology), 20ng/mL EGF (Sigma), 10μg/mL Heparin (StemCell Technologies), 10μg/mL Insulin (Sigma), and 1μg/mL hydrocortisone (Sigma). Three wells per population were plated for each animal (3 animals per genotype). Spheres were imaged at 200x and counted on day 9. Mammosphere size was measured from the captured images.

#### 3D Colony/Organoid Forming Assay

After 9 days of suspension culture, 50 mammospheres (from above) per sample replicate were resuspended into 60uL of Matrigel in a 96-well ultra-low attachment plate. The sphere-Matrigel drop was allowed to solidify inside a 37°C, 5% CO_2_ incubator for 15 min before covering with MMS medium supplemented with 5% FBS. Media was refreshed every 5 days. After 9 days organoids were counted and imaged at 100x.

#### 3-Dimensional Culture of Cells on Matrigel

Cells were plated into 8-well chamber slides at a density of 2×10^4^ cells/well in 100uL GFR-Matrigel. After Matrigel solidification, overlay 3D MEC media was added (phenol red free DMEM/F12, 5% horse serum, 20ng/mL EGF, 10ug/mL Heparin, 20ng/mL bFGF, 0.5ug/mL hydrocortisone, and 10ug/mL insulin). Cells were fed with fresh media every 3 days until day 10 when cells were imaged or processed for immunofluorescence.

#### Generation of *NF1* mutant MCF10A cells using CRISPR

Target-specific single guides for *NF1* were designed to match the rat-specific guides from Dischinger *et al*..^20^ Guide 1 (g1; 5’ GGTCCAGTCAGTGAACGTAA 3’) and guide 2 (g2; 5’ AATAGCATTGGATACAGAGC 3’) both were targeted to human *NF1* in exon 21. LentiCRISPR v2 was a gift from Feng Zhang (Addgene plasmid # 52961; http://n2t.net/addgene:52961; RRID:Addgene_52961). The annealed DNA was cloned into the bsmb1 sites. The plasmid was co-transfected with packaging plasmids (CMV-VSV-G and psPAX2 from Addgene) into HLA 293T cells to package the lentivirus. Collected lentiviral supernatants were added to MCF10A cells and were treated with puromycin until non-transfected control MCF10A cells died. The resultant *NF1* mutant lines were maintained as pooled populations because individual cell cloning was unsuccessful.

#### RNA-seq

RNAseq was performed on MCF10A EV and *NF1* mutant cell lines using 3 biological replicates. 1.2×10^5^ cells were plated into one well of a six-well plate. 24 hours later, buffer RLT (from RNeasy kit) containing betamercaptoethanol was added to each well of cells. The mix of buffer RLT and cells was then added to an MPbio lysing matrix E tube. Cells were homogenized for 20 seconds in an MPbio homogenizer, then the supernatant was added to a QIAshredder column. An equal volume of 70% ethanol was added to the flow through, and the entire volume of liquid added to an RNeasy column. RNA was extracted according to the manufacturer’s instructions.

Libraries were prepared by the Van Andel Genomics Core from 500 ng of total RNA using the KAPA mRNA Hyperprep kit (v4.17) (Kapa Biosystems, Wilmington, MA USA). RNA was sheared to 300-400 bp. Prior to PCR amplification, cDNA fragments were ligated to IDT for Illumina TruSeq UD Indexed adapters (Illumina Inc, San Diego CA, USA). Quality and quantity of the finished libraries were assessed using a combination of Agilent DNA High Sensitivity chip (Agilent Technologies, Inc.), QuantiFluor^®^ dsDNA System (Promega Corp., Madison, WI, USA), and Kapa Illumina Library Quantification qPCR assays (Kapa Biosystems). Individually indexed libraries were pooled and 100 bp, paired-end sequencing was performed on an Illumina NovaSeq6000 sequencer using an S4, 200 cycle sequencing kit (Illumina Inc., San Diego, CA, USA) to an average depth of 70M reads per sample. Base calling was done by Illumina RTA3 and output of NCS was demultiplexed and converted to FastQ format with Illumina Bcl2fastq v1.9.0

Demultiplexed FASTQ files were trimmed of sequencing adapters using Trim Galore v0.6.0 (https://www.bioinformatics.babraham.ac.uk/projects/trim_galore/) and then mapped to GRCh38 with GENCODE release 33 with STAR v2.7.8a basic two-pass mode.^69^ The raw counts were extracted from the reverse strand alignments. PCA was computed in R using the normalized log counts from edgeR and the prcomp function from the stats package. Using the normalized counts per million, the fold change against average EV expression was calculated for all samples and shown as log2(fold change) capped at −5 and 5. Otherwise, color indicates Z-scores across all displayed samples. GSEA was done in R using the msigdbr^70^ and clusterProfiler^71^ packages with 10,000 permutations and genes ranked by fold change.

### QUANTIFICATION AND STATISTICAL ANALYSIS

Automated segmentation and quantitation of epithelial cells from surrounding stromal cells was performed using Aperio Slide Manager software (12.4.3.5008) on the Aperio AT2 (Leica Biosystems). Fold change in epithelial cell percentage relative to wildtype was plotted with ggplot2 (v3.3.3) in R (v4.0.3). A beta regression was used to determine if there were significant differences between wildtype and mutants using the betareg (v3.1-4) package and emmeans (v1.5.4) package for Tukey adjusted contrasts.

For bud counts, total alveolar and TEBs were counted in a blinded manner by three individuals in four animals per genotype. A Poisson generalized linear mixed-effects model that included random intercepts to account for multiple scorers and repeated animal measurements was generated using the lme4 package (v1.1-26) in R (v4.0.3). Emmeans (v1.5.4), with Tukey adjusted p-values, was used to determine if there were significant differences in bud count between the wildtype and mutants and results were plotted with ggplot2 (v3.3.3).

For colony counts, a negative binomial generalized linear mixed-effects model that included a random intercept to account for repeated animal measurements and an unstructured covariance matrix was used to determine if there were significant differences in the number of colonies formed between the wildtype and mutant rats for the luminal and basal populations using the lme4 (v1.1-26)- package and the emmeans (v1.5.4) package for Tukey adjusted contrasts in R (v4.0.3).

Mammosphere count and size data were stratified by luminal and basal populations. A negative binomial generalized linear mixed-effects model was used to determine if there were significant differences in mammosphere count between the wildtype and mutant rats. For mammosphere size, data were log transformed and a linear mixed-effects model was used to determine if there were significant differences in mammosphere size between the wildtype and mutant animals. Both models included a random intercept to account for repeated animal measurements, an unstructured covariance matrix, and Tukey adjustment of the p-values. The R (v4.0.3) packages lme4 (v1.1-26) and emmeans (v1.5.4) were used.

For 3D counts, a negative binomial generalized linear mixed-effects model that included a random intercept to account for repeated animal measurements and an unstructured covariance matrix, was used to determine if there were significant differences in the number of 3D organoids formed between the wildtype and mutant rats using the lme4 (v1.1-26) package and emmeans with a Tukey adjustment (v1.5.4) package in R (v4.0.3).

For the CytoFlex analysis of 21 vs 43 day old rats, beta mixed-effects model with a random intercept for each animal were used. There was significant evidence for several three-way interactions: viability x cell-type x genotype, timepoint x viability x cell-type, as well as a timepoint x cell-type x genotype (p = 0.008, p < 0.0001, and p < 0.0001 respectively). Therefore the final model included these three three-way interactions, their composite two-way interactions, and individual main-effects. Hypotheses of interest were assessed via linear contrasts with two-sided hypothesis tests, Benjamini-Hochberg adjusted p-values, and targeted a <5% false discovery rate. Results are reported with Tukey 95% confidence intervals.

## ACKNOWLEDGMENTS

We would like to thank the VAI Vivarium staff, Flow Cytometry Core, Genomics Core, and Optical Imaging Core for their valued expertise. Thank you to all the memes that made this paper possible. This study was supported by the U.S. Department of Defense Neurofibromatosis Research Program (W81XWH-21-1-0759), the Breast Cancer Research Foundation, Bee Brave Foundation, Muskegon Tempting Tables, NF Michigan, and the Van Andel Foundation.

## Data and code availability

The raw data of bulk RNA-seq generated by this study have been deposited in NCBI’s Gene Expression Omnibus and are accessible through GEO Series accession number GEO: GSE209545. Accession numbers of each dataset are listed in the key resources table. This paper does not report original code. Any additional information required to reanalyze the data reported in this work paper is available from the lead contact upon request.

## Author contributions

Conceptualization, C.R.G, M.R.S, E.A.T.; Methodology, C.R.G, E.A.T., P.S.D; Validation, C.R.G., E.A.T.; Investigation, E.A.T., M.A., C.J.E, P.S.D, M.C., R.S., L.T., C.E.; Formal Analysis, J.L.G., Z.B.M., I.B., K.F.; Writing – Original Draft, E.A.T., C.R.G; Writing – Review & Editing, E.A.T, C.R.G, M.R.S.; Funding Acquisition, M.R.S, C.R.G; Resources, Supervision, M.R.S., C.R.G.

