## Supplemental Figures for "*Nf1* deficiency accelerates mammary development and promotes luminal-basal plasticity"

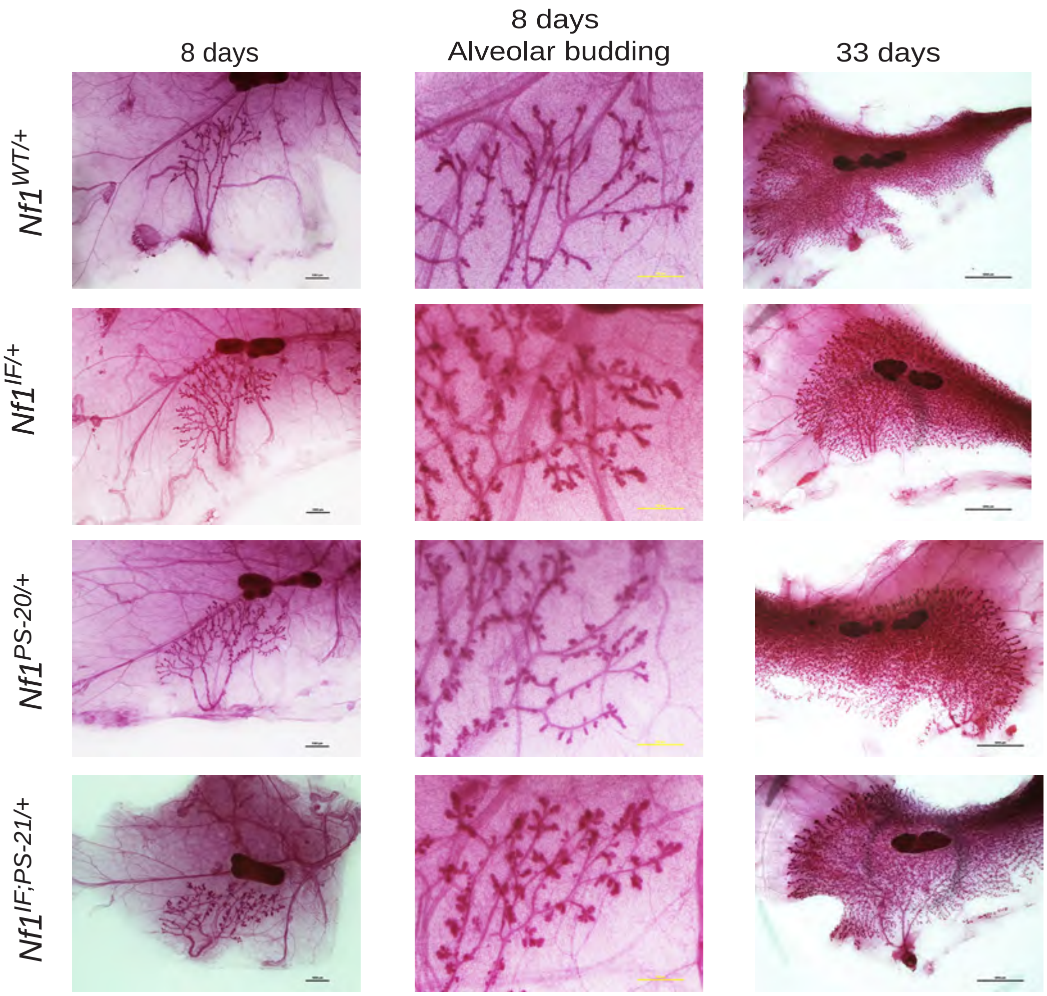


**Figure S1: Accelerated ductal outgrowth and aberrant alveolar budding is present at PND 8 in *Nf1*-deficient mammary glands.** Carmine stains of whole mammary fat pads from wild-type and *Nf1*-deficient rats showing the entire ductal tree in eight-day old females at 20x (first column, scale bar = 1000 µm), ductal branches at 80x in eight-day old females (second column, scale bar = 500 µm), and the ductal tree in 33 day old females (last column, scale bar = 5000 µm) at 8x magnification.

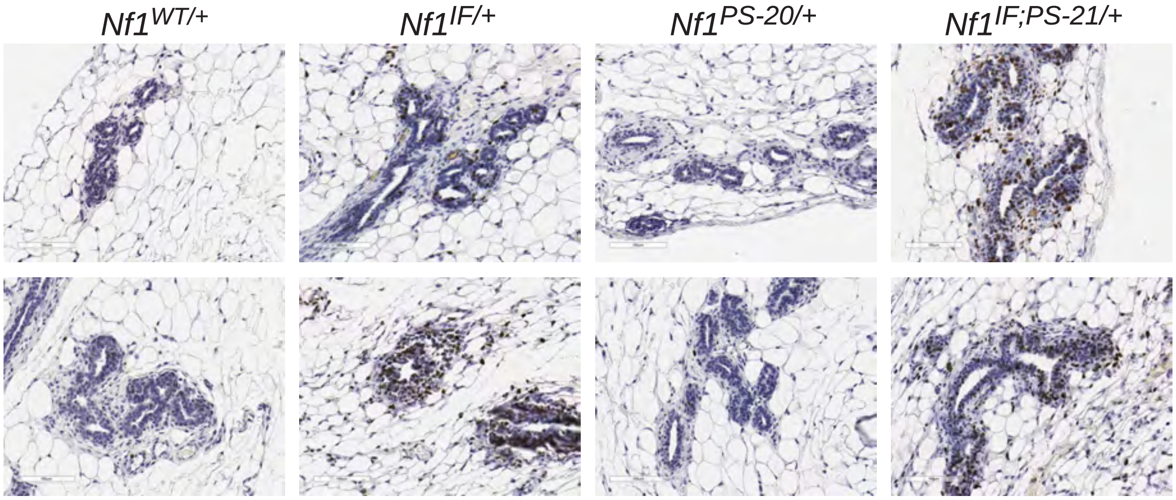


**Figure S2: Increased proliferation observed in *Nf1*-deficient mammary glands.** Ki67 immunohistochemistry of mammary duct sections of 21-day old female rats. Images were taken at 200x, scale bar = 100 µm.

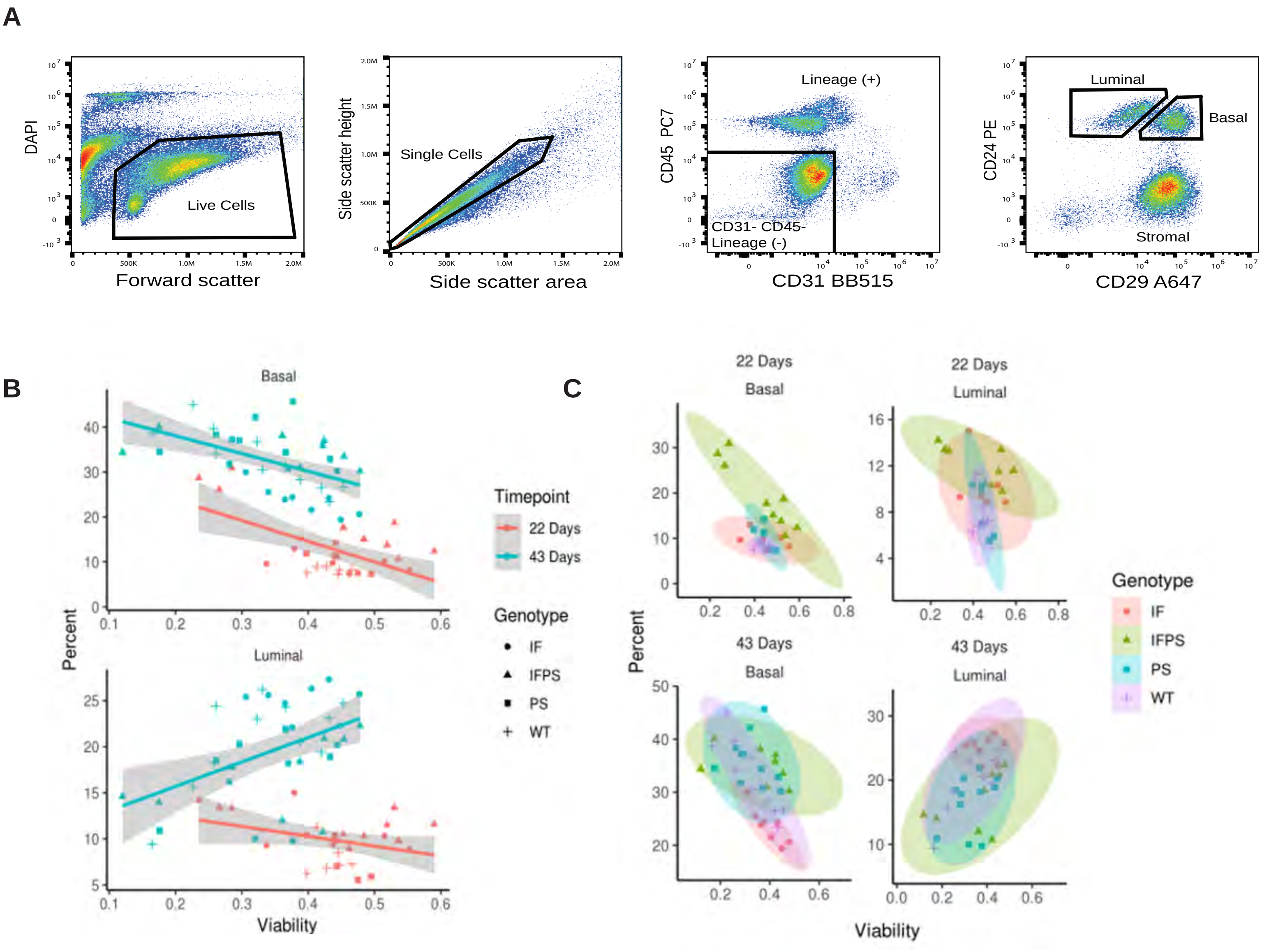


**Figure S3: Luminal/basal cell sorting and analysis of cell viability, timepoint, and genotype interactions.** A) Flow cytometry gating strategy for the isolation of CD31-CD45-CD24+CD29+ mammary epithelial cells. B&C) The effect of flow sorted cell viability on luminal or basal population percentage varied by timepoint, cell-type, and genotype.

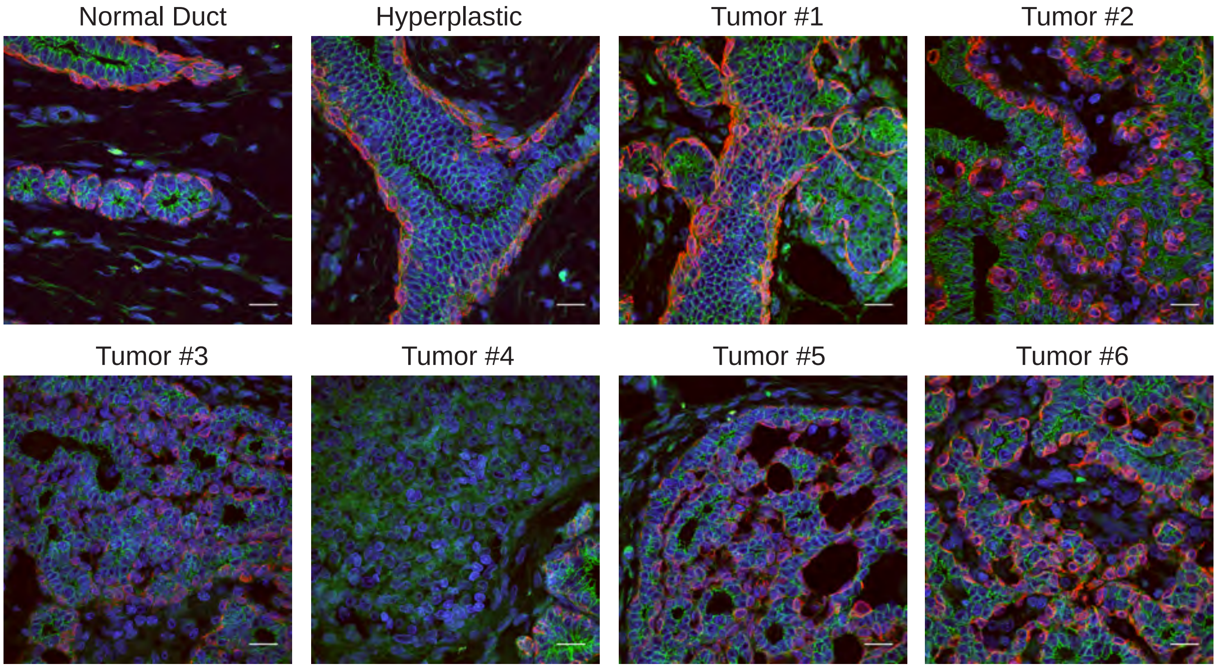


**Figure S4: Presence of Ecad+/CK14+ cells in normal, hyperplastic, and tumor tissue from a single *Nf1*-deficient mammary gland.** Confocal images from a single tissue section of a normal mammary duct, hyperplastic duct, and tumors from a *Nf1*$^{PS\text{-}20/+}$ female. The tissue was stained with CK14 (red), E-cadherin (green), and DAPI (blue) and imaged at 600x magnification (scale bar = 21 µm).

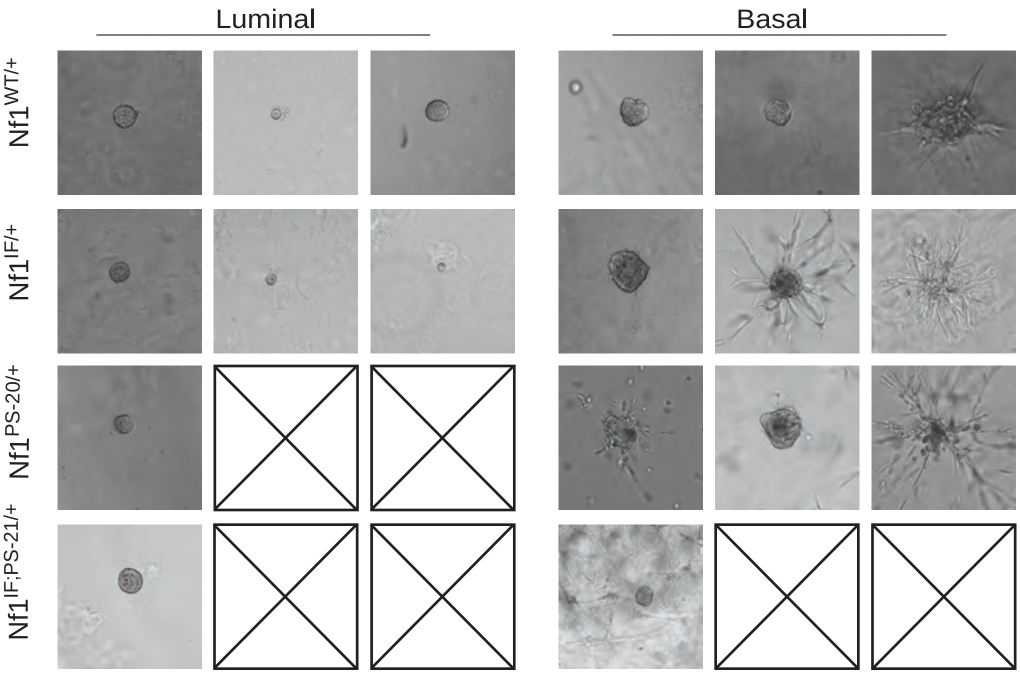


**Figure S5: Luminal *Nf1$^{PS\text{-}20/+}$* and *Nf1$^{IF;PS\text{-}21/+}$* cells are unable to form 3D colonies.** Representative images of 3D organoids formed from 50 mammospheres (from part C) embedded in Matrigel. Images were taken at 100x magnification, scale bar = 400 µm. Boxes with an x indicate samples where none of the replicates formed organoids.

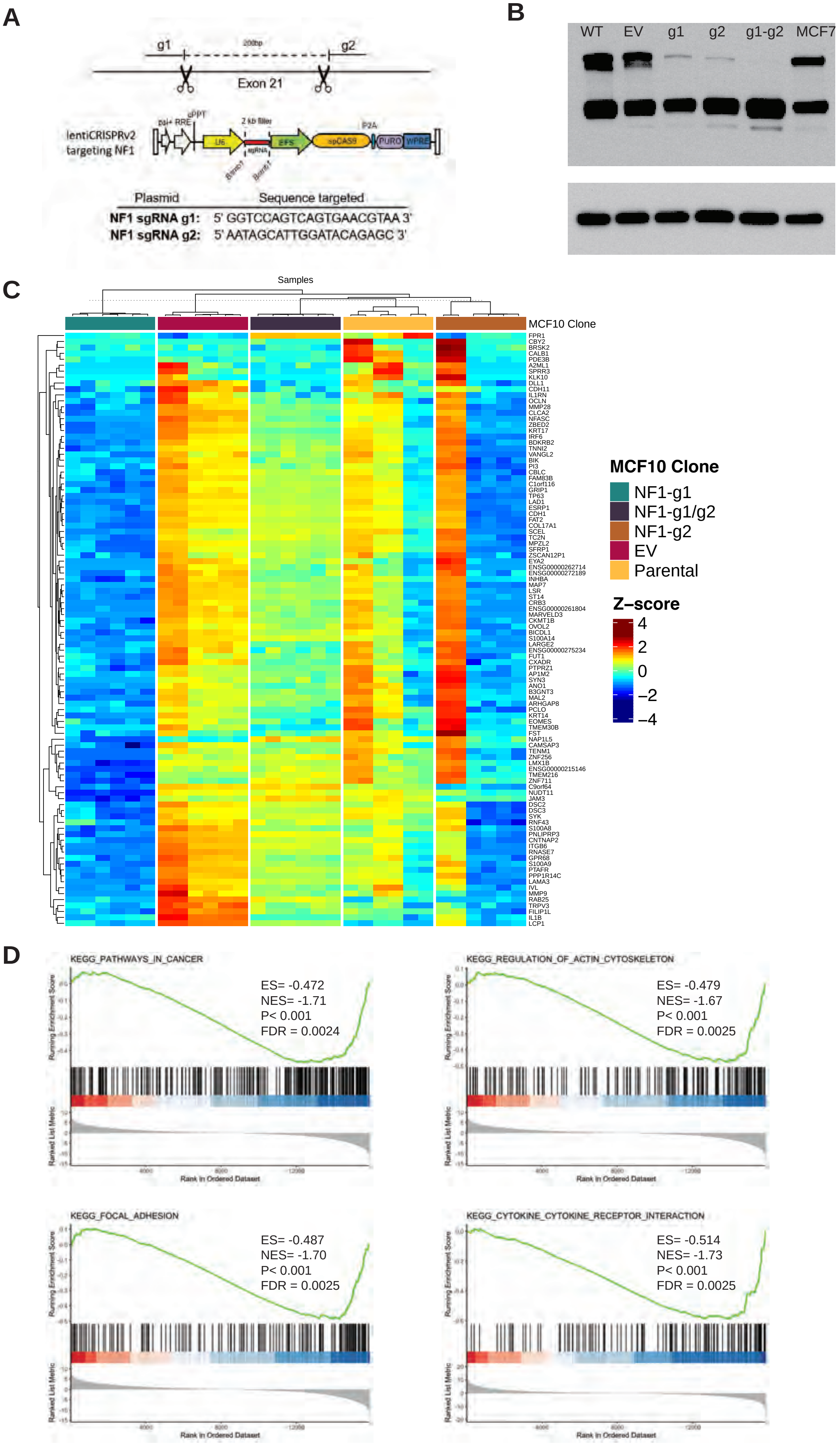


**Figure S6: MCF10A *NF1*$^{mut}$ cell lines have altered signatures related to cancer progression.** A) Schematic showing CRISPR guides targeting exon 21 of the human *NF1* gene. B) Immunoblot analysis of neurofibromin in *NF1* CRISPR mutants compared to empty vector, parental MCF10A and MCF7 cells. C) Heatmap of the top 100 differentially expressed genes in the *NF1* CRISPR mutants, empty vector, and parental MCF10A cells from RNA sequencing analysis. D) KEGG pathways enriched in the *NF1* CRISPR mutants from GSEA of RNA sequencing data.
